# Planar aggregation of the influenza viral fusion peptide alters membrane structure and hydration, promoting poration

**DOI:** 10.1101/2021.08.09.455689

**Authors:** Amy Rice, Sourav Haldar, Eric Wang, Paul S. Blank, Sergey A. Akimov, Timur R. Galimzyanov, Richard W. Pastor, Joshua Zimmerberg

**Author notes:** Division of Virus Research and Therapeutics, CSIR-Central Drug Research Institute, Lucknow 226031 (UP), India. A.R. and S.H. contributed equally to this work.

## Abstract

To infect, enveloped viruses employ spike protein, spearheaded by its amphipathic fusion peptide (FP), that upon activation extends out from the viral surface to embed into the target cellular membrane. Here we report that synthesized influenza virus FP are membrane active, generating pores in giant unilamellar vesicles (GUV), and thus potentially explain both influenza virus’ hemolytic activity and the liposome poration seen in cryo-electron tomography. Experimentally, FP were heterogeneously distributed on the GUV at the time of poration. Consistent with this heterogeneous distribution, molecular dynamics (MD) simulations of asymmetric bilayers with different numbers of FP in one leaflet show FP aggregation. At the center of FP aggregates, a profound change in the membrane structure results in thinning, higher water permeability, and curvature. Ultimately, a hybrid bilayer nanodomain can form with one lipidic leaflet and one peptidic leaflet. Membrane elastic theory predicts a reduced barrier to water pore formation when even a dimer of FP thins the membrane as above, and the FP of that dimer tilts, to continue the leaflet bending initiated by the hydrophobic mismatch between the FP dimer and the surrounding lipid.

## Introduction

For all enveloped viruses, one or more glycoproteins on the surface of the viral membrane mediates the fusion of the envelope to the cell membrane for transport of the viral genome to the target cell cytoplasm, bringing about infection. In the first electron microscopy visualizations of purified viral spike proteins from rabies, rubella, influenza, and other viruses, a striking similarity between spikes from different viruses was the assembly of these purified viral spike proteins into aggregates, termed rosettes, as their hydrophobic trans-membrane domains (TMDs) aggregated ^1-8^. Most enveloped viruses enter their target cells via the endocytic pathway, where the viral envelope spike protein encounters acidic pH. For the influenza virus spike protein hemagglutinin (HA), acidic pH activation of HA is necessary and sufficient for triggering fusion of the viral envelope to a variety of target membranes, including receptor-doped phospholipid bilayers ^1,2,9-15^. In the absence of their TMDs, activation of isolated soluble ectodomains of HA led anew to fresh rosettes of those trimers ^1,2^. The N-terminal domain of HA2 is responsible for this second aggregation of HA ectodomains – it is a short amphiphilic N-terminal sequence that became known as the fusion peptide (FP). As with the first rosette of HA, this second rosette formation is considered a consequence of the hydrophobic effect: the hydrophobic surface formed by one side of the FP would avoid water via association with the hydrophobic surface of another FP. The influenza FP comprises the N-terminal 21 amino acids of HA2, located within the HA ectodomain (at neutral pH), proximal to the HA trimer surface but near the TMD. At low pH the FP is found in the target membrane, as evidenced by hydrophobic photolabeling ^16,17^. The FP is required for infection in vivo and membrane fusion in vitro and is featured in all hypotheses on HA-mediated fusion, though there is little agreement on the structural, compositional, and mechanistic data to date on its exact role ^18,19^. Since the FP is a highly conserved region of the influenza virus genome across many different subtypes of influenza virus ^20,21^, and a universal feature of enveloped viral fusion proteins, determining the FP’s role for infectivity and membrane fusion is critical to finding variant-independent immunogens and pan-viral therapeutics to ameliorate morbidity and mortality.

We recently studied influenza hemifusion intermediates in vitro with high-resolution cryoelectron tomography using a phase plate to enhance signal-to-noise by a factor of four ^22^. The cryotomograms of influenza viral like particles and receptor-laden liposomes showed the expected 10 nm diameter hemifusion diaphragm in high cholesterol-containing target liposomes, but at lower cholesterol, exposed membrane edges were detected on the target membrane in direct contact with one or more HA still attached at their transmembrane base to the viral envelope; this lipid-protein structure can extend the FP outward from the virus linearly for many nanometers. These surprisingly stable large ruptures in the membrane are reminiscent of some of the earliest and most consistent clinical findings in virology (hemolysis) and *a)* suggest that fusion under these conditions is target-leaky to even large macromolecules and *b)* are consistent with the loss of target liposome contents in studies of intact virus-liposome fusion and a similar cryo-electron tomography study ^23,24^. The target membrane lipid dependence for poration is also recapitulated for intact virions in a single vesicle dye entry assay ^25^. From the existing literature on vesicle-vesicle fusion, it is hard to determine if the influenza FP alone is responsible for bilayer poration since earlier studies collected data during concurrent lipid mixing and phase changes ^26-28^. The purpose of this study is to determine whether, and how, the influenza FP by itself could initiate stable pores in lipid bilayers.

Here, in a study of the membrane mechanisms by which influenza virus can disrupt a target membrane, we establish that FPs underly this disruption: in target membranes, a reversible pore forms upon addition of FP in the absence of virus or even the rest of HA. In MD simulations crafted to understand the chemistry by which FP act, a third kind of rosette emerged: the aggregation of FP via their lateral side chains (not their hydrophobic surfaces) into FP microdomains that displace lipids in the cis leaflet. This novel structure locally thinned the bilayer and significantly increased the probability of water entry. A new model is proposed to explain our data based on a tilting of FPs towards each other to further thin the remaining lipids immediately under even an FP dimer. For larger aggregates, this more hydrated, thinner membrane structure replaces the lipid bilayer in a small domain wherein a lipidic pore can form.

## Results

### Experimental Results: Fusion peptide is sufficient to porate GUV

Stable pore formation, detected as passive transport of the water-soluble, membrane impermeable fluorescent dye Alexa 488 into (influx) or out (efflux) of GUV, was monitored by confocal microscopy. *Influx*: GUV are immersed in Alexa 488 containing solution. Prior to FP addition, the interiors of GUV were dark. Upon addition of fluorescently labeled FP at neutral pH, FP bound to GUV (Fig. 1A, red). In fractions of the GUV (here defined as the leakage fraction), Alexa 488 accumulated inside the GUV changing the black interior to green (Fig. 1A, Influx). In other fractions of GUV, leakage was not detected and the absence of poration was deduced despite the binding of FP: no Alexa 488 influx was detected (Fig. 1A No Influx). The fluorescence signal increased until the signals detected inside (F_in_) and outside (F_out_) of the GUV were equal (Fig. 1B; representative examples; n=7 from 2 independent GUV preparations); the influx ratio *R* = *Fin*/*Fout* = 1 is defined as 100% influx. GUV integrity was maintained during and after poration (the intensity and distribution of DiD lipid dye, labeling the GUV membrane, remained unchanged). *Ef f l ux*: Poration was observed as a loss of GUV internal fluorescence with the efflux ratio, following normalization, *R* = (*Fin* – *Fout*)/(*F*_*avg*_(t=0)-*Fout*) = 0 defined as 100% efflux (data not shown). POPC GUV encapsulating 3,000 MW Alexa 488-labeled dextran showed 100% efflux into a non-fluorescent bathing solution (n = 2 from 1 GUV preparation). FP induced pores in POPC GUV were a) stable on time scales that allow equilibration of the probe with the external medium, and b) large enough to allow transport of both 3,000 and 570.5 MW molecules. Thus, the poration of GUV seen upon acidification of attached influenza virus or viral-like particles (VLP) was recapitulated with the addition of one part of the influenza HA -- the FP.

**Fig. 1.**
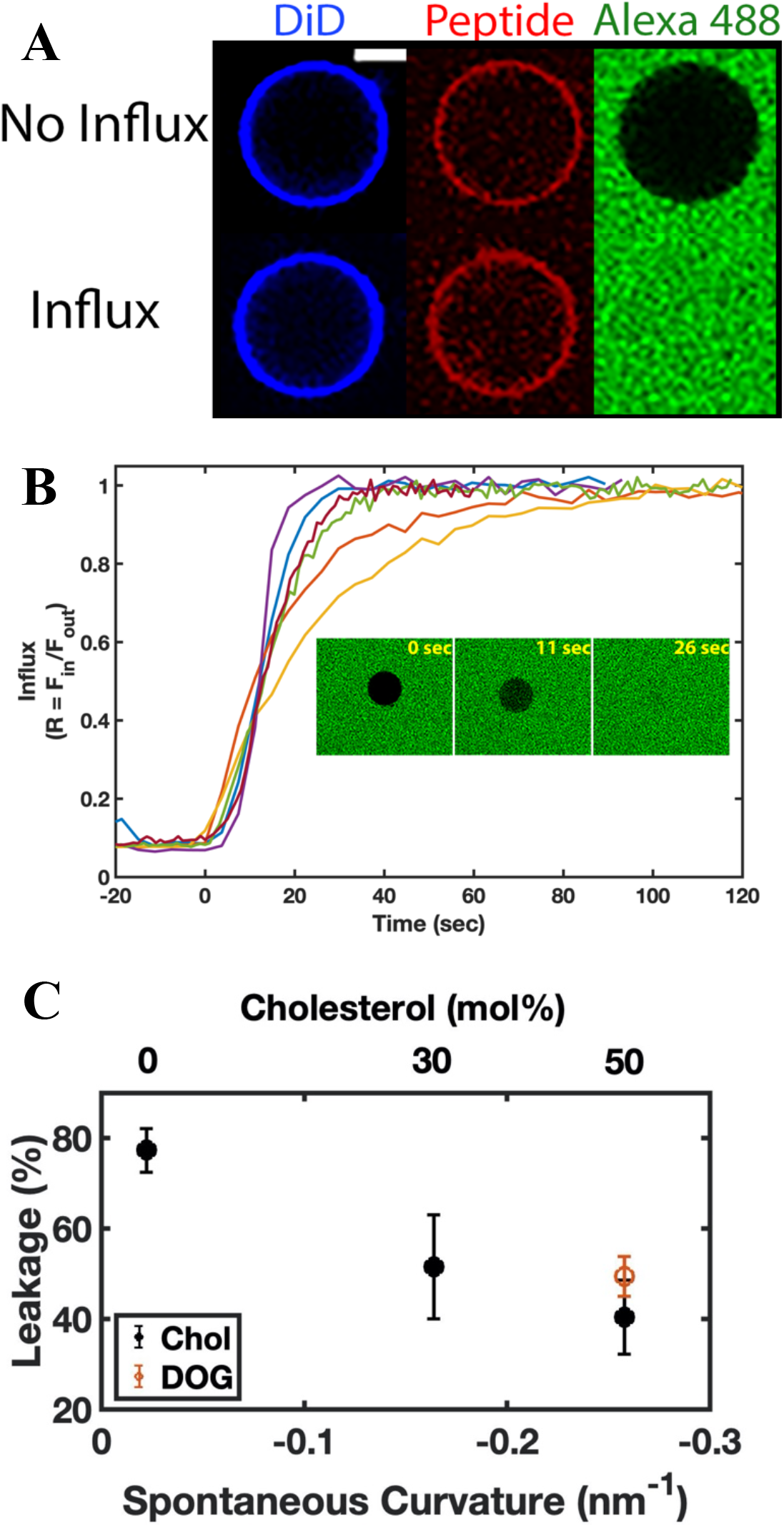
Influenza fusion peptide induced poration of GUV. (A) Representative examples of DiD labeled GUV exhibiting binding of the influenza FP and no flux (top) or influx (bottom) of Alexa 488 containing solution following FP addition. (B) Kinetics of influx following FP addition. The ratio of Alexa 488 average intensities in the GUV lumen and the external solution as a function of time. Examples from 6 different experiments. For clarity, differently colored data are aligned in time such that 0 time indicates the onset of intravesicular fluorescence significantly above background. Inset shows a typical example of vescle undergoing influx at 11 and 26 sec after onset of influx (0 sec). White scale bar 5 µm. (C) Fusion peptide induced poration (leakage) is dependent on target membrane lipid composition. Error bars represent the SEM of three independent preparations or the range of 2 (30% cholesterol). Closed circles: 0, 30, and 50 mol% cholesterol in POPC. Open red circle: 24 mol% DOG in POPC.

Since viral-induced pores were preferentially detected in target membranes containing low cholesterol (< ∼40 mol %) ^22,25^, the dependence of FP-induced pores on lipid composition was tested. Target membranes comprising higher cholesterol concentrations should hinder FP-induced poration. In the presence of FP, vesicle poration (Leakage %) of POPC GUV with varying cholesterol concentrations decreased with increasing cholesterol concentration, consistent with our hypothesis (Fig. 1C, n=3 independent GUV preparations evaluating 50 - 100 GUV per experiment); leakage decreased from ∼80% to 40% for cholesterol concentrations 0 to 50 mol% corresponding to leakage decreases of 34% and 48% relative to 0 mol% cholesterol.

The ring structure of cholesterol confers rigidity to membranes and this property could explain the observed decreasing leakage with increasing cholesterol. To determine whether cholesterol induced rigidity or if negative monolayer spontaneous curvature (MSC) of the GUV leaflet is a correlating parameter, GUV were prepared with the same MSC (∼ – 0.258 nm^−1^) as those of the highest cholesterol concentration used in this study (50 mol %), by replacing cholesterol with 24 mol % dioleoyl glycerol (DOG). The MSC of cholesterol, DOG, and POPC used in the calculations of the mixture MSC are: –0.494 nm^−1^, –0.99 nm^−1^, and –0.022 nm^−1^, respectively ^29,30^ and we use the commonly accepted approximation that the MSC of a lipid mixture is defined as a concentration weighted average of the component MSCs ^28^. 24 mol % DOG GUV did not exhibit statistically different leakage fractions from 50 % cholesterol GUV (40.3 ± 8.2 % for 50 mol % cholesterol compared to 49.3 ± 4.4 % for 24 mol % DOG; mean± SEM, *n* = 3, respectively; *p* = 0.39, 2-tailed equal variance T-test; Fig. 1C). These results are consistent with the hypothesis that leakage and MSC are correlated parameters, and lipids with negative MSC, e.g. cholesterol or DOG, hinder pore formation ^31^.

According to the classical theory ^31^, lipids can be described by their effective “molecular shapes”: (*i*) conical lipids have larger tail than headgroup cross-sectional areas, which leads to negative MSC; (*ii*) inverted conical lipids have smaller tail than headgroup cross-sectional areas leading to positive MSC; and (*iii*) cylindrical lipids have zero MSC. Lipids with positive MSC (inverted cone) tend to form structures with positive geometrical curvature like the pore edge, thus enhancing membrane poration and inhibiting fusion. Lipids with negative MSC (cone) tend to form structures with negative geometrical curvature like fusion sites, thus inhibiting membrane poration and enhancing fusion.

To gain insight into the relative lifetime of the FP induced pores with and without cholesterol, pore stability was monitored via the influx ratio, R, in the presence and absence of 50% cholesterol (Supplementary Fig. S1A). Pure POPC vesicles exhibited *R* close to 1 (> 80%, *n* = 14 of 17 reported, have filling ratios > 0.95), consistent with stable pores over a time sufficient for equilibration with the outside concentration to occur (tens of seconds, Fig. 1B). In the presence of cholesterol, *R* ranged from ∼0.1 to 0.6 (> 90%, *n* = 28 of 30 reported, have R < 0.6). The sub-maximal level of influx in these GUV and estimates of pore open time (Supplementary Fig. S1B) are consistent with a higher probability of pore closing in the presence of high cholesterol ^32^. Thus, FP induced pore lifetime is shorter in cholesterol/POPC GUV than in POPC GUV.

To further confirm whether submaximal leakage indicates a pore closing event or a slowly flickering pore, a second marker (sulforhodamine B, SRB) was introduced 25 minutes later following the introduction of the first marker, Alexa 488 ^33^. Since SRB and Alexa 488 have similar molecular weights (559 and 643 g/mol, respectively), if a pore allows entry of Alexa 488 then the same pore should allow influx of SRB. Consistent with this hypothesis, when introduced simultaneously, Alexa 488 and SRB have comparable levels of influx (Supplementary Fig. S2A). However, if an initially open pore closes, then the influx of SRB would be restricted. Supplementary Fig. S2B shows the influx of Alexa 488 (the first soluble marker) when incubated in the presence of the FP, but, in this example, SRB was excluded when introduced later, i.e., there was no influx. Hence, over this time scale, FP induced pores are not stable in the presence of 50% cholesterol although they are stable in membranes without cholesterol.

Lastly, the question of FP aggregation was considered. Analyses of the distributions of FP in bulk solution and on the vesicle indicated that, in addition to an increase in the average amount of FP on the vesicle in time, the variability at the pixel scale (∼500 nm^2^) also increases in time. In all experiments, the distribution of pixels for the best fit intensity surface indicated the presence of pixels with larger intensity deviations compared to solution; the differences between the peptide in solution and on the GUV is not explained by the differences in intensity but is consistent with the appearance, in time, of a non-uniform or segregated distribution of fusion FPs. Specifically, poration occurs with a characteristic time, 101 +/-22 sec, and normalized FP density, 2.7 +/-0.05, (mean +/-SEM; n = 9 including both Alexa 488 and dextran experiments), both log-normally distributed, where the characteristic time represents the difference between the time leakage is first detected and the time the normalized FP fluorescence on the vesicles increases above the FP fluorescence in solution (see Fig. 9 and aggregation analyses in Methods). The combined deviations from n = 9 experiments taken around the estimated poration time are shown in Fig. 2. The intensity deviations for FP on the GUV were broader with extended tails, relative to the solution distribution, indicating that many pixels have higher than expected intensities under the null hypothesis that the distribution of FP on the vesicle is the same as the solution distribution. The difference in dispersion between the two distributions is significant (n = 16,200 for both distributions, alpha = 0.00001, Ansari-Bradley test). This observation is consistent with aggregation of FP at the optical pixel scale. The pixels with higher-than-expected peptide intensities are ∼ 10 times greater than the average intensity in solution. With a nominal voxel size of 1×1×2 µm^3^ and all peptides in the voxel contributing to the intensity measured in a nominal 1×1 µm^2^ pixel, a rough estimate of the number of peptides present at these “hot spots” is ∼ 36,000. Assuming a uniform distribution at the level of a single voxel, the number of FPs, when scaled to the MD simulation size (15 nm x 15 nm) is ∼ 8; a value comparable to that used in MD simulations.

**Fig. 2:**
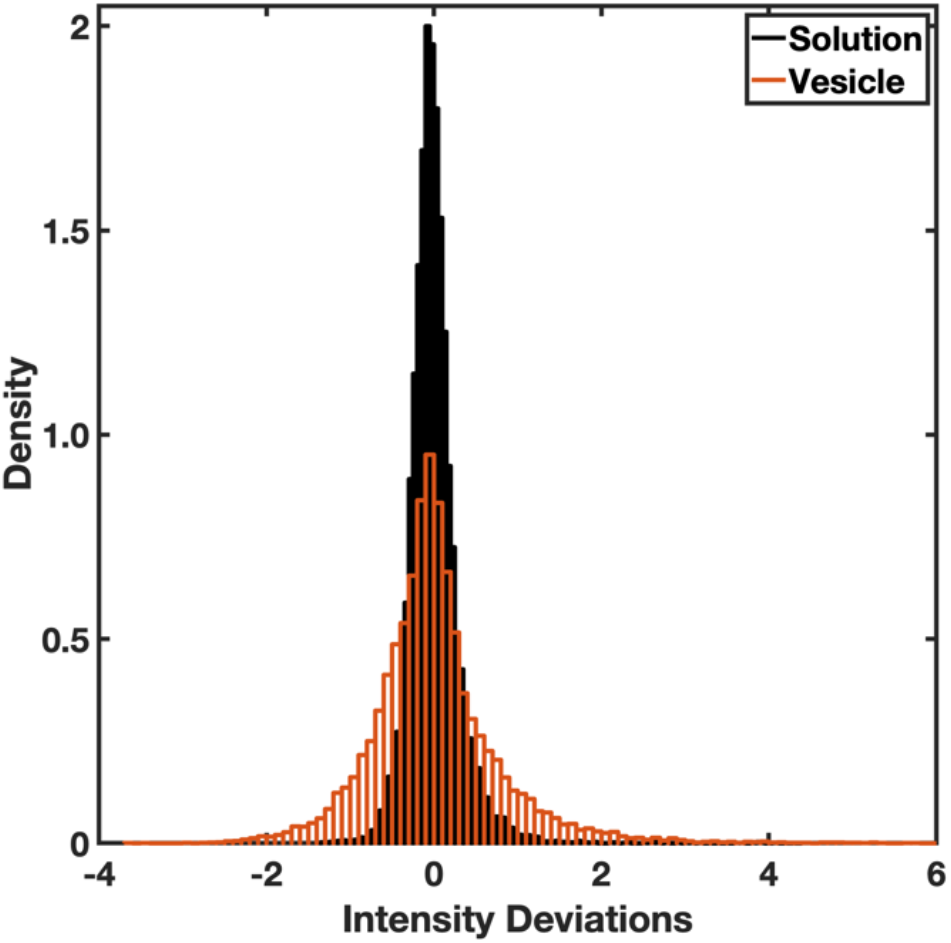
Peptides on vesicles at the estimated poration are not distributed randomly. The intensity deviations in solution (black) are narrower around 0 deviation (mean = -3.9e-5, variance = 0.096) while the intensity deviations for the bound peptide (red) include deviations greater than the largest deviations observed in solution (mean = 0.005, variance = 0.598).

The GUV experiments support the hypothesis that the FP moiety of influenza HA, when aggregated, is sufficient to porate membranes in the absence of the entire protein structure. The poration process is dependent upon GUV lipid composition. To explore the role of membrane composition, FP density, and potential FP interactions (peptide-peptide and peptide-lipid) in the poration process, molecular dynamics simulations of asymmetric membranes with varying numbers of FP were evaluated at the 1:1 chol:POPC bilayer composition that gave the maximal inhibition of poration.

### Molecular Dynamics Simulations

Simulations were performed with 1 (2.1 μs), 6 (2.1 μs), or 10 (21 μs) FP interacting with one leaflet of either POPC or chol:POPC membranes (Fig. 3). While the top-down images of 6 FP show little aggregation, there are clearly significant interactions among peptides for the 10 FP systems. These interactions are correlated with a loss of lipid tails beneath the clusters in the 10 FP systems for both POPC and chol:POPC and are a consistent feature observed for all cluster sizes detected throughout their trajectories (Fig. 3). Fig. 4 shows top-down views of both 10 FP systems at 0, 11, and 21 μs as well as the cluster compositions as a function of time; Fig S3 presents the number and compositions of clusters for each microsecond. As these figures show, the peptides are separated at the start of the trajectory, and dimers are formed in the first several μs. In POPC, most dimers are relatively short-lived, lasting 2 μs or less, though a dimer of FP 7-9 remains stable from 5-21 μs (the end of the trajectory). There is qualitatively more clustering in chol:POPC, with 4 long-lived dimers (FP 8-10, 7-8, 1-9, and 5-7); these form shorter-lived clusters of as many as 6 FP with each other and assorted monomers. Hence, there is considerable mixing over 21 μs in both systems. Of note, the most stable dimers are antiparallel, with the two N-terminal helices interacting with one another. Interactions between FPs are primarily mediated by direct FP–FP interactions, rather than water or ion bridges. This aggregation may be surprising given the net negative charge of these FPs. However, because even a single FP leads to local membrane deformations, clustering of FPs decreases the net energy of the FP-lipid boundary of the system which stabilizes clusters (see Theory section below). Aggregation in the 6 FP systems is much more limited (Figure S4), as consistent with the lower peptide concentration. A detailed examination of aggregation as a function of concentration will be reported in future work using coarse-grained models.

**Fig. 3.**
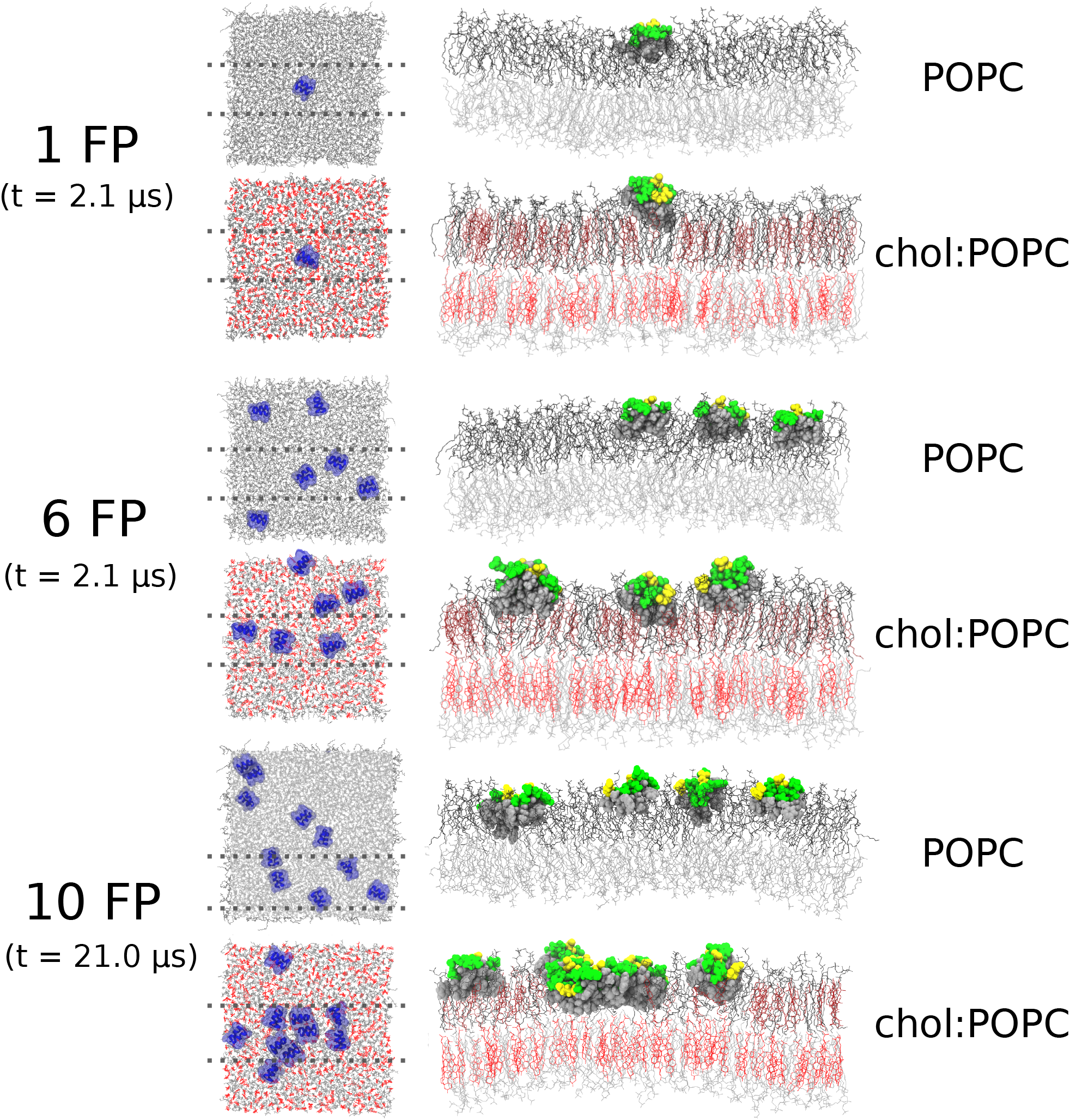
Snapshots of microsecond time-scale MD simulations. Final simulation snapshots. POPC and cholesterol are shown in grey and red line representations; FPs are depicted as blue-ribbon diagrams with a translucent space-filling overlay in the top-down views. Side view of the final simulation snapshots are taken as a 40 Å section denoted by the dotted lines in the top-down views; lipids in the cis leaflet are depicted in darker shades to distinguish them. FPs are depicted in a Van der Waals sphere representation, with hydrophobic residues in grey, while polar residues are green and acidic residues are yellow.

**Fig. 4.**
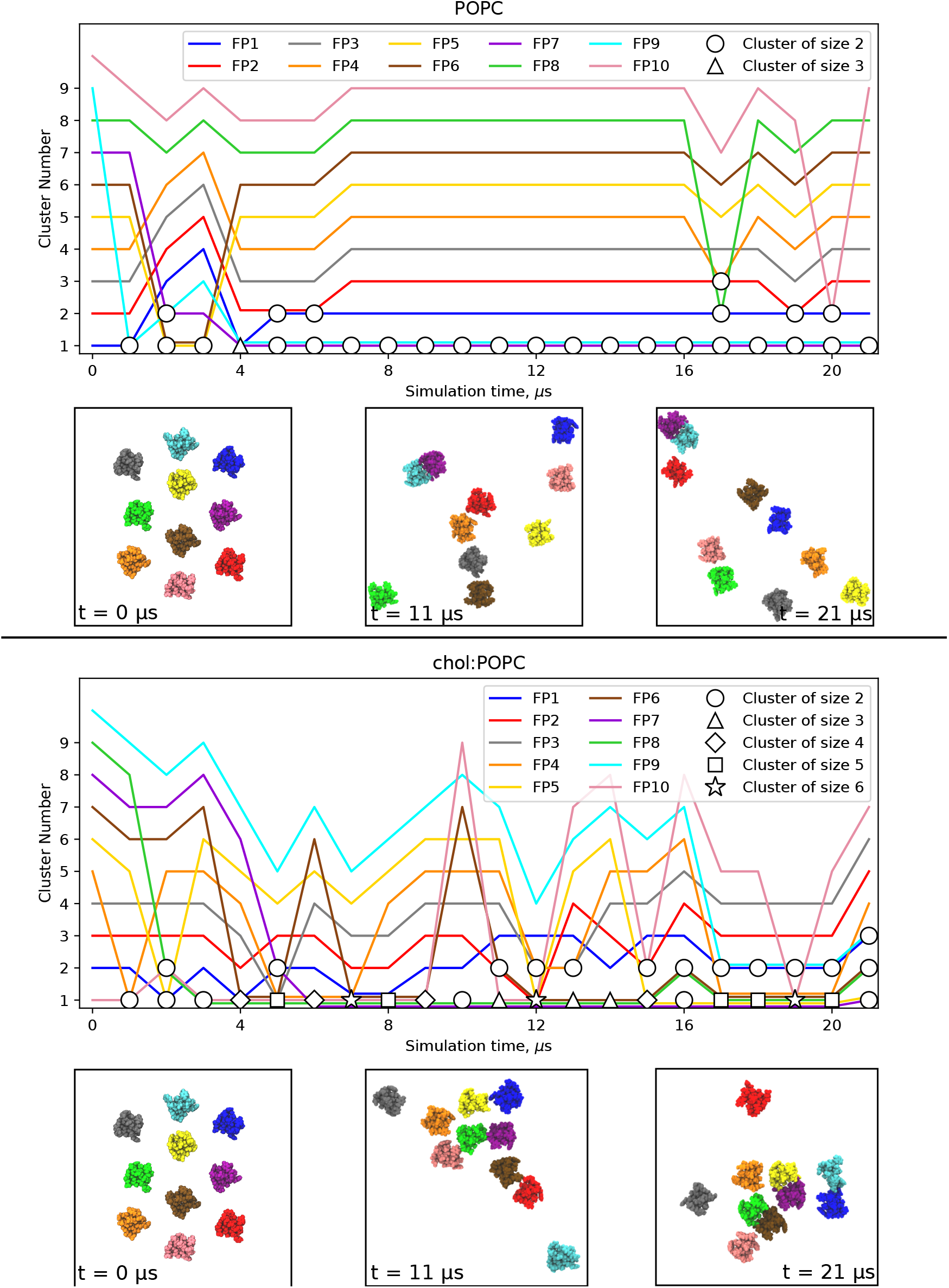
Snapshots and clustering of microsecond time-scale MD simulations with 10 FP. Snapshots are shown at times 0, 11 and 21 μs of the 21 μs simulation. Plots show the cluster composition at each microsecond, with the symbol representing the number of peptides in a given cluster. The peptide colors in the plot correspond to the colors in the snapshots.

The effect of the FP clustering on the membrane is profound: aggregates displace lipids in the upper (cis) leaflet, curve the trans leaflet (Figs. 3 and S5), and sit much lower in the leaflet than an isolated FP (Fig. S6). As shown in Fig. 5A for two snapshots of FP in POPC from the 10 FP simulation, even an antiparallel dimer can displace lipids, and thereby come in direct contact with the trans leaflet. This can be understood as follows: FP monomers are “cradled” by adjacent lipids, as has been observed previously for extended surface associated amphipathic helices^47^. In contrast, the inserted area of the dimers is too large to be covered by lipid in the same leaflet, and consequently the lipids are replaced by the peptide aggregate. The top two panels of Fig 5B show views of the underside of the entire cis leaflet at t=0 for both 10 FP bilayers. The hydrophobic undersides of the FP are only faintly visible indicating negligible lipid displacement. The bottom two panels of this figure show substantial displacement of lipids by 3 dimers in POPC at 17 µs, and by a hexamer in chol:POPC at 19 µs.

**Fig. 5.**
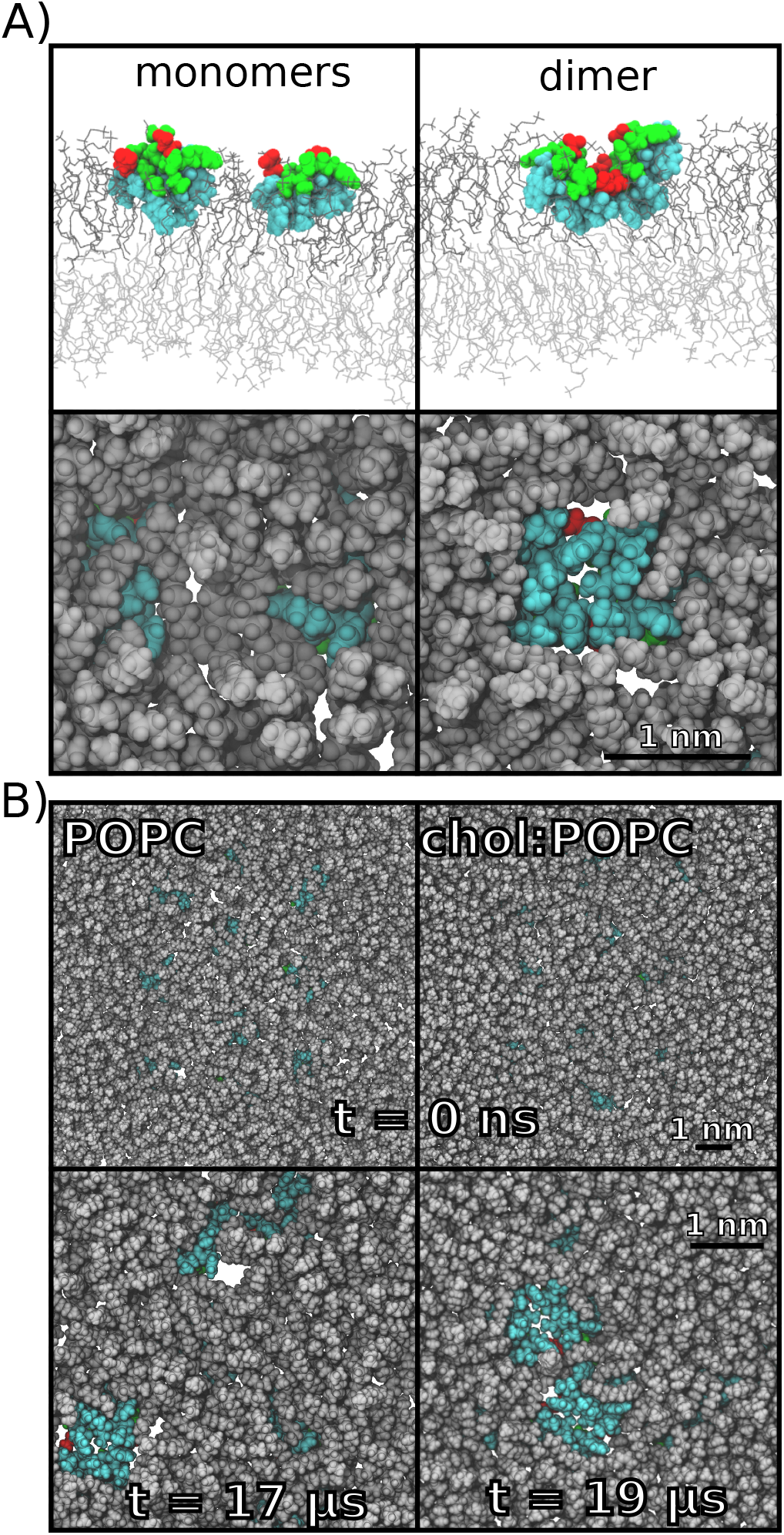
Dimers displace lipids. A) Side view (top panels) and bottom view (bottom panels) of two FP in POPC when they are isolated (left, t = 1 μs) or in an antiparallel dimer (right, t = 15 μs). Peptides are shown as Van der Waals spheres with hydrophobic residues in cyan, acidic residues in red, and polar residues in green. In the side view, top leaflet lipids are depicted as dark grey lines while bottom leaflet lipids are in lighter grey. In the bottom-up view, only lipids in the top leaflet are shown, and they are depicted as grey spheres. B) Same view as above, but for the entire POPC (left) and chol:POPC (right) leaflets at t = 0 when the peptides are isolated. As peptides cluster over the course of the simulations, lipid displacement becomes more pronounced. Lipid displacement was observed whenever clustering occurred. The undersides of the three dimers at 17 μs are shown for POPC (left), and for the hexamer at 19 μs in chol:POPC (right); the FP identities can be read from Fig. 4. Black scale bars are 1 nm in all cases.

The structure of the helical hairpin also promotes dimerization and insertion. Specifically, irrespective of clustering or insertion depth, all FP in all simulations were seen as “rotated” from a reference state in which both N- and C-terminal helical components of the FP would be the same average distance from the bilayer midplane. This rotation was quantified by a roll angle ρ, (Fig. S8), with the N-terminal helix of the hairpin beneath the C-terminal helix; similar rotations were reported by Brice and Lazaridis from simulations of a single FP in DMPC ^34^. The FP roll angle was larger in the chol:POPC membranes, with <ρ> =56.0° ± 0.7 as compared to 51.6° ± 0.8 in pure POPC (mean ± SEM; p < 0.005, pooled t-test). The long axis of the peptides remained roughly parallel to the membrane surface, regardless of aggregation or membrane composition, though large (> 25°) positive or negative tilt angles are occasionally sampled (Fig. S8, tilt angle).

This orientation of the FP differs from that of most amphipathic peptides, which consist of a single long alpha helix lying relatively flat on the bilayer surface. Partial vertical stacking of the two FP helices increases the peptides’ thickness projection along the bilayer normal, with the FP subtending a total thickness of ∼15 Å. However, the rolled orientation of the FP does not prevent water molecules from penetrating this sparse layer, unlike the lipid leaflet which they replace. This rolling and concomitant thickening along the z-axis allow the FP assembly to replace lipids in the cis leaflet more easily than a linear amphipathic peptide would, and with less hydrophobic mismatch at the cluster boundary. Consistent with the increased leaflet thickness of chol:POPC compared to pure POPC, the larger value of ρ allows the FP to span a longer distance along the z-axis. Movies of each of the system snapshots shown in Fig. S3, demonstrating peptide tilt and roll, are included in the Supporting Materials.

Lipid displacement by the FP aggregate in turn leads to localized membrane thinning (Fig. 6 upper left). Thinning is most pronounced in the 10 FP chol:POPC system, which has the densest FP aggregates. The chol:POPC:10FP system thins by 4.5 Å in the region of the aggregate, to a bilayer hydrophobic thickness of 31.0 Å compared to 34.5 Å in the peptide-free case (Fig. 6). The POPC:10FP system thins by a more modest 2 Å (from 27.5 Å to 25.4 Å). This combination of thinning and lipid displacement leads to substantial deformation of the trans leaflet (Figs. S3 and S5), as the leaflet bows towards the cis leaflet, so that the terminal methyl groups of the trans leaflet lipids can contact the hydrophobic underside of the FP aggregate. As anticipated, the extent of this deformation becomes more pronounced with increasing aggregate size (Fig. S5). Because chol:POPC is a thicker bilayer than POPC and displays a greater extent of FP-generated thinning, the explicit curvature of the trans leaflet is more prominent in these systems (Fig. S5). Also, the peptide rides higher into the aqueous media bathing the cis leaflet in chol:POPC when surrounded by lipid but inserts deeper into the membrane to abut the trans methyl groups, so the thinning itself is greater in absolute terms, and the lipid bounding the FP has a greater deformation in chol:POPC. In the theory on FP aggregation (below) this increased deformation creates a large line tension around the FP cluster and explains the slight bias towards FP dimer formation seen in chol:POPC (Figs. 3 and S3).

**Fig. 6.**
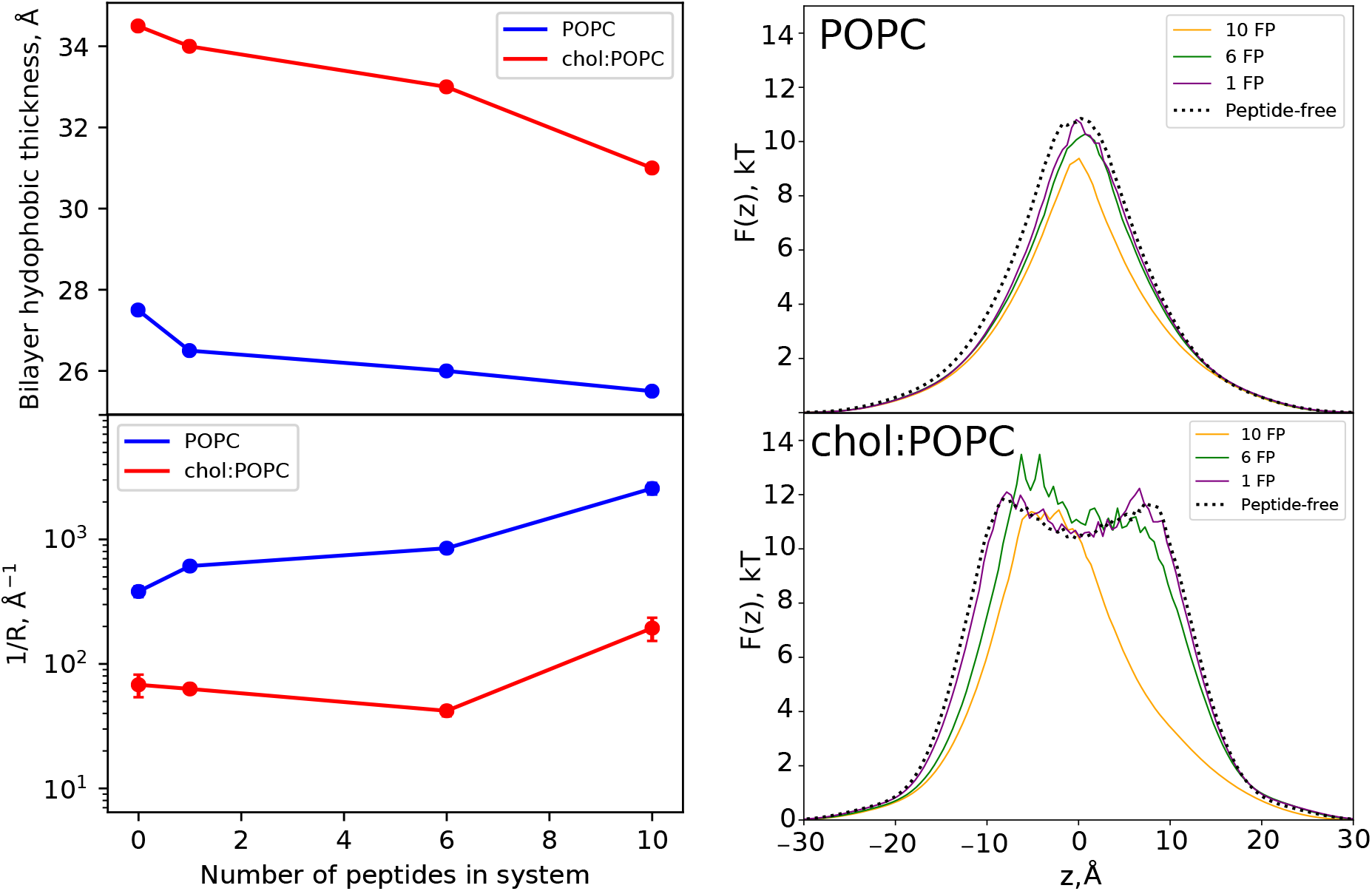
Thickness and water properties for all systems simulated. Only atoms contained in a cylinder of radius 30 Å and centered at the origin are included (see Fig. S7). FPs are in the cis leaflet (z > 0). Left Top) hydrophobic thickness of the POPC and chol:POPC membrane vs numbers of FP. Left Bottom) Values of inverse resistances to water permeability 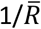 (Eq. 7) vs numbers of FP. Error is reported as standard error of the mean, calculated over 250 ns block averages for all systems, except for the peptide-free cases where 100 ns block averages were used. Error bars are smaller than the symbols for some systems. Right) Free energy *F*(*z*) (Eq. 6) for water as a function of position along the membrane normal for pure POPC

These geometric changes modulate the spontaneous curvature of each leaflet in ways not well-described by the traditional theoretical model of non-interacting FPs. The FP aggregate-dependent displacement of lipids in the cis leaflet, and the high degree of localized bilayer thinning, suggest that bilayers with FP present would be more susceptible to pore formation.

While the conventional MD simulations presented above are not of sufficient length to generate pores or pre-pore configurations, they do yield information on water content. The free energies, *F*(*z*), for water in the FP-containing and FP-free POPC and chol:POPC bilayer systems are plotted in Fig. 6 (right) across the thickness of the bilayer. These profiles are obtained from the probability distributions of water (eq. 6 in Methods) and are commonly termed potentials of mean force (PMF). All systems show low free energy in the well-hydrated headgroup region and a substantial free energy barrier in the hydrophobic hydrocarbon region. The barrier at the midplane (*z* = 0) for pure POPC is characteristic of most homogenous bilayers ^35^; the metastable minimum at the midplane in chol:POPC results from chain ordering and is found in liquid ordered phases ^36^. Addition of a single FP only slightly lowers the free energy barrier, while the aggregates observed in the 10 FP simulations substantially alter the energy landscape. The barrier height is reduced by approximately 1.6 k_B_T for POPC:10FP, and the width at half-height by roughly 3 Å; the barrier height is relatively unchanged for chol:POPC:10FP but the width at half-height is reduced by 12 Å, indicating significant membrane thinning.

Furthermore, the plateau region of the PMF around z=0, which is characteristic of cholesterol-containing membranes, is absent in the 10 FP system. Aggregates of intermediate sizes from the 6 FP simulations have an analogous effect following the same trend as the extreme cases of 10 FP and FP-free – reduction of the barrier height and concomitant narrowing in the POPC membranes and narrowing with an associated shape change in chol:POPC. Since pore formation is presumed to be initiated by the formation of a water wire (see Theory section below), these reductions in the free energy for water can be taken as an indirect measure of the propensity of the aggregates to generate pores. The formation of hydrophilic pores is developed in the following section.

The free energy of water in each membrane can be related to the permeability by the inhomogeneous solubility diffusion (ISD) model (eq. 7) ^37^. Fig. 6 (lower left) plots the inverse resistances to water permeability 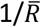 evaluated from *F*(*z*) and the ISD model (eq. 8). Since the permeability *P* is proportional to 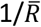, it is most instructive to compare values of 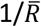 between systems. The presence of FP aggregates leads to a ∼7-fold increase in 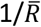 for the case of 10 FP in pure POPC – while the 10 FP aggregate has a more moderate ∼3-fold increase for the chol:POPC membrane. Even the presence of a single FP increases this ratio from ∼5 to 10, indicating a stronger effect on the pure POPC membrane. Most notably, 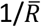for POPC:10FP is 13-fold larger than for chol:POPC:10FP, implying that the permeability to water is lower by a factor of 13 when cholesterol is present.

### Theory

#### FP clustering

The inhomogeneous distribution of FPs on the surface of GUV at the time of poration is consistent with the hypothesis that, following FP surface binding, FP aggregation is an important driver of poration. The MD simulations at varying FP concentrations show a cooperative activity of multiple FPs: aggregates thin the membrane and exclude cis leaflet lipid tails (Fig. 6) and illustrate elastic deformation of the membrane at the FP/membrane interface for even a single FP. Thus, FPs incorporated into the lipid leaflet can be considered as generators of boundary conditions that induce membrane deformations. We calculated the elastic contribution to the FP cluster boundary energy in the framework of the theory of lipid membrane elasticity ^38^ assuming symmetric FP, as described in Material and Methods.

Bringing two FPs into close contact eliminates part of the peptide/membrane boundary, and consequently nullifies the deformations induced by that part of the boundary. Thus, nonzero boundary energy acts as a driving force for aggregation of FPs. Entropy-based forces lead to the lateral dispersion of the FPs at lower overall concentrations. The balance of these two forces results in some critical concentration of FPs, above which aggregation becomes possible. To ensure aggregation, the total free energy difference Δ*E* between the ensemble of clusters containing *n*_*clust*_ FPs each and the system of separated FPs should be negative. We combine the elastic energy of the cluster (*E*_*cluster*_), the elastic energy induced by single FP (*E*_*FP*_) and the free energy arising from the entropy of mixing (*E*_*ent*_) into Δ*E* = *E*_*cluster*_ – *E*_*FP*_ + *E*_*ent*_ and require Δ*E* to be negative as a condition for spontaneous clustering. FP aggregation is ensured when the normalized surface concentration (*x*) of FPs is above a critical concentration of FP (*x*_*c*_) where *x*_*c*_ is the value of *x* satisfying the equation Δ*E*(*x*) = 0. For large clusters (*nclust* ≫ 1) one can neglect the boundary energy of the cluster and entropy of the cluster ensemble (see Appendix in SI). The approximate expression 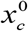 for the critical concentration reduces to a simple Boltzmann factor of the energy induced by insertion of a single FP (for details see SI):

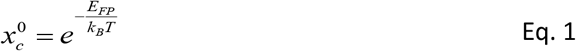

For the elastic parameters specified in the Material and Methods (*Calculations of the energy of elastic deformations of lipid membranes*.), 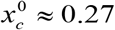. The dependence of the critical concentration *x*_*c*_ for aggregation on cluster size is weak and demonstrates that Eq. 1 is a reasonable estimate (see Fig. S13). The optimal cluster size is calculated from the total cluster formation energy. There is a clear energy minimum at the cluster size ∼10 (Fig. 7). Note that this minimum is local, and at larger sizes the energy again decreases, predicting that larger systems should form macroscopic clusters. Thus, small FP clusters can form spontaneously, while the formation of larger clusters requires overcoming an energy barrier. Accounting for dipole-dipole attraction, one finds the formation of hydrogen bonds between contacting FPs will decrease the critical concentration *x*_*c*_ and enhance the formation of the cluster.

**Fig. 7.**
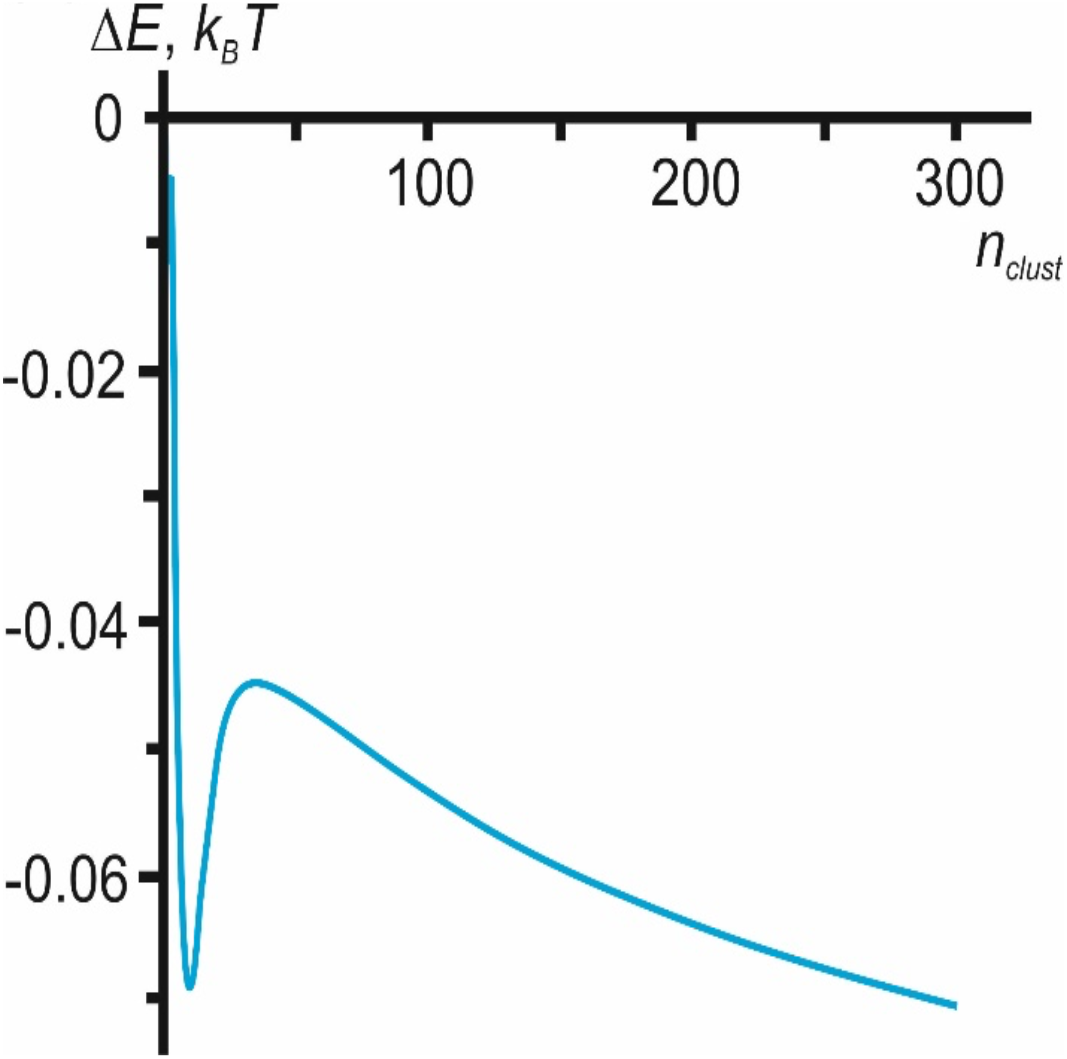
Dependence of cluster formation energy on cluster size (per 1 FP). Dependence of the energy of cluster formation on its size for the super optimal FP concentration *x* = 0.3. Additional contributions to clustering are related to dipole-dipole interactions and hydrogen bonds between contacting FPs, but they are not accounted for in these calculations.

#### The pathway to pore formation

Since a cylindrical aqueous pore in a bilayer is the best pathway for lipid pore formation, the increased hydration of the FP domain bilayer (Fig. 8c, right) should lower the energy of formation of a critical intermediate state, the hydrophobic defect, without lipid reorientation. ^39^ Here the side walls of the water filled defect is formed by hydrophobic lipid tails. The surface tension at the interface of lipid tails and water depends non-linearly (although monotonically) on cylinder radius, as water structure inside a small hydrophobic cavity differs from that in bulk water. This difference in water structure is accounted for in the framework of Marčelja theory^40,41^, where the energy of a homogeneous hydrophobic cylinder filled with water is:

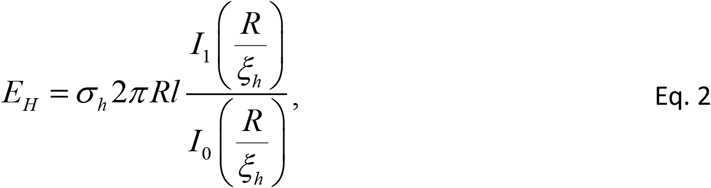

where *σ*_*h*_ is the surface tension at the macroscopic interface of lipid tails and water, *ξ*_*h*_ ∼ 1 nm is the characteristic length of hydrophobic interactions, and *I*_1,0_ are modified Bessel functions of the first and zeroth order, respectively. According to Marčelja’s theory, hydrophobic surface–water interactions are described by an order parameter that characterizes the hydrophobic surface induced perturbation of the water structure. The associated free energy density is decomposed into a series with respect to the order parameter scalar field and its gradient. The final energy of the hydrophobic surface exposition is calculated by minimizing the total energy functional with respect to the order parameter distribution. The dependence of the cylindrical hydrophobic defect energy on its length *l* is linear. The length of the cylinder, *l*, and the cylinder radius, *R*, completely describe the state of a membrane pore, and with these two parameters, one can build continuous trajectories – from intact bilayer (*R* = 0, *l* = 2*h*, where *h* is the leaflet thickness), to the hydrophilic pore (*l* = 0, *R* > 0). These trajectories have common features: 1) the transition of the hydrophobic defect to a subsequent hydrophilic pore requiring the surmounting of an energy barrier, and 2) the energy of the hydrophilic pore has a local minimum at some radius (approximately equal to the leaflet thickness *h*), i.e., the hydrophilic pore can be metastable ^39^. The metastability is provided by the fact that, at a pore radius around typical lipid leaflet thicknesses, meridional positive curvature cancels equatorial negative curvature, making this region less stressed ^39^. In POPC bilayers, the energy barrier to pore formation is high enough (∼40 *k*_*B*_*T, k*_*B*_*T* ≈ 4 × 10^−21^ J) to essentially prohibit spontaneous pore formation. This agrees with the experimental observation (Fig. 1) that the dye leakage from the GUVs occurs only after FP application. We previously found that the maximum of this energy barrier occurs at a hydrophobic defect radius of *R* ≈ 0.7 nm; the critical defect radius is weakly dependent on the elastic parameters of the membrane ^39^.

**Fig. 8.**
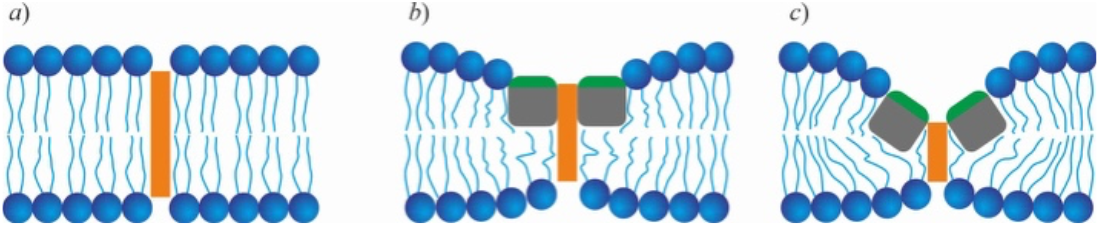
Three possible ways to form a hydrophobic defect (orange rectangle). a) purely lipid membrane; b) flat FP cluster; c) funnel-like structure in the FP cluster formed by tilted FPs. The hydrophobic defect length decreases from *a* to *c*.

There are two main ways to induce membrane destabilization: (*i*) decreasing the energy of the hydrophobic defect by membrane thinning and (*ii*) making the pore edge energetically preferable by the addition of components with positive MSC, e.g. lysolipids. The membrane-disrupting amphipathic peptide piscidin 1^42^ acts in the first way. Once embedded in the cis lipid leaflet, it can tilt to form a funnel-like structure that locally decreases bilayer hydrophobic thickness, promoting hydrophobic defects. Our MD simulations showed that FPs decrease membrane thickness (Figs. S3 and 5) which should facilitate formation of hydrophobic defects (compare Figs. 7a and 7b). However, the average bilayer thinning induced by FPs and the thickening induced by cholesterol are comparable: a chol:POPC:FP membrane has approximately the same thickness as a pure POPC membrane, and thus both should have comparable energy barriers to pore formation. In contradistinction, experimentally the former membrane is leaky, while the latter is almost non-leaky indicating an additional effect of FP apart from average membrane thinning occurs. This contradiction can be reconciled if FP clustering and tilting of the FPs towards each other in the cluster (Fig. 8c) leads to formation of a funnel-like structure. Now, the hydrophobic region of the *cis* leaflet almost vanishes, and the opening of the pore only requires a hydrophobic defect to span the *trans* leaflet.

We calculated the energy barrier towards pore formation for each system considered in the MD simulations. The barrier is defined by the energy of the water cylinder spanning the bilayer (*E*_*hydro*_) and the deformation of the membrane near the hydrophobic defect (*E*_*def*_). The energy is evaluated relative to the energy of the state of distantly separated (minimally interacting), *n* FPs (*nE*_*FP*_).

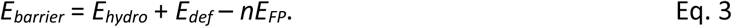

To calculate the membrane deformation energy, we utilized the Hamm & Kozlov ^38^ elastic model essentially in the same way used to calculate the deformation energy arising at the FP cluster boundary. We assume that the system is axially symmetric with respect to the axis passing through the center of the FP cluster or single FP and introduce a cylindrical coordinate system, *O-r-φ*. As FPs are hypothesized to tilt, specific boundary conditions were imposed onto the projections of lipid directors (time-averaged lipid tail orientations) onto the *O-r* axis (*n*_*r*_(*r* = *R*, φ)) and *O-φ* axis (*n*_*φ*_ (*r* = *R*, φ)) of a cylindrical coordinate system. The boundary director depends on the FP tilt angle, FP insertion depth, and membrane thickness. Three reference points are considered for determining the dependence of the boundary director on these parameters. The first one is the single FP; the *n*_*r*_-projection of the boundary director (*n*_*mono*_) is defined from geometrical considerations ^43,44^:

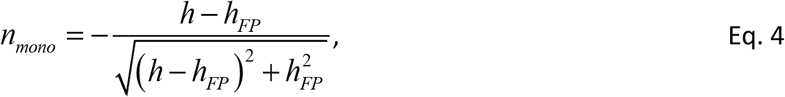

where *h*_*FP*_ is FP hydrophobic thickness. By symmetry, the *n*_*φ*_−projection of the boundary director is zero. The second reference point is the membrane, squeezed at the FP boundary to match its hydrophobic thickness. Now the boundary director *n*_*h*_ is defined from the condition of the deformational energy minimum. The last reference point is the FP cluster, where boundary peptides are tilted inwards or outwards of the cluster, and the hydrophobic and leaflet thicknesses match. In that case, the boundary lipid director equals the FPs tilt *θ*, describing the rotation of the FP’s long axis (see Fig. S8). We interpolate these three reference points linearly:

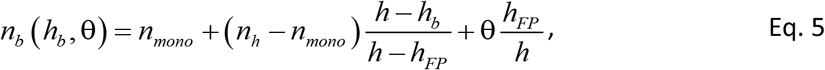

where *h*_*b*_ is the thickness of the lipid leaflet at the point of contact with FP. The energy of the cylindrical hydrophobic defect is calculated in the framework of the Marcelja model ^40,41^ according to Eq. 2 ^39^. The calculation procedure was as follows. For a purely lipid membrane, we accounted for both the hydrophobic and elastic deformation energies arising in the hydrophobic defect vicinity. For FP-containing lipid membranes, we calculated the energy of membrane deformation around the FP cluster, in which FPs are tilted inwards by a 45° angle. The hydrophobic defect in this case spans only the *trans*-leaflet. The lipid boundary director was set in accordance with Eq. 5; hydrophobic defect length was set equal to the hydrophobic thickness of the *trans*-leaflet. The total energy was minimized with respect to the membrane deformations and the depth of the FP insertion *h*_*b*_ (Fig. 8c). The hydrophobic defect radius, *r* = 0.7 nm, corresponding to the maximum of the energy barrier of pore formation, weakly depends on the elastic parameters ^39^ and was taken equal for each system under consideration amended by the DOG-containing system. We assumed that the latter system has MSC equal to the cholesterol-containing one (as in the GUV experiments). The thickness and rigidity of this membrane are taken equal to the POPC membrane, as unsaturated DOG lipid tails should not affect lipid ordering. The results of the calculation are summarized in Table 1. There are two drivers of the poration inhibition – MSC and thickness of the membrane leaflet. A negative MSC restrict FP tilting, while increased thickness raises the energy of the hydrophobic defect. The chol:POPC system has the highest energy barrier for pore formation due to the highest bilayer hydrophobic thickness. The effect of FPs on the membrane poration is explained by the cooperative action of several FPs that can form a cluster capable of thinning the membrane during a fluctuation of mutual orientation and thus decreasing the energy barrier towards pore formation. DOG and cholesterol, due to their approximately equal negative MSC, inhibit pore formation in the presence of FPs by impeding FP mutual tilting. This suggests that it is not primarily the chemical structure of cholesterol *per se* that is responsible for cholesterol’s pore neutralization activity. However, cholesterol’s ability for leaflet ordering can inhibit membrane poration additionally by increasing membrane rigidity. An increase in FP concentration leads to an increased probability of cluster formation, and the larger cluster further lowers the barrier to pore formation; both factors facilitate pore formation.

**Table 1.** Barriers to pore formation depend on lipid composition and FP cluster size.

| Mixture | Pore barrier, $k_B T$<br>(FP cluster = 4) | Pore barrier, $k_B T$<br>(FP cluster= 2) |
| --- | --- | --- |
| POPC | 40.4 | 40.4 |
| POPC:FP | 14.7 | 19.1 |
| chol:POPC | 53.3 | 53.3 |
| chol:POPC:FP | 22.5 | 25.5 |
| DOG:POPC | 42.7 | 42.7 |
| DOG:POPC:FP | 27.6 | 25.9 |

## Discussion

For viral pathogenesis, the essential topological event in cell entry occurs when viral and host cell membranes merge, an activity mediated by the FP. Here, synthetic HA FPs segregate on lipid bilayers to high density in occasional ‘hot spots’ at the time that pores form, replicating intact influenza virion and viral-like particles in porating POPC and cholesterol:POPC lipid bilayer membranes ^25^. This HA FP-induced poration was inhibited, and poration lifetime reduced by including cholesterol in the target membrane. To investigate mechanisms, in silico MD simulations were implemented on lipid bilayers with increasing FP densities. Surprisingly, a novel structure emerged at higher FP densities: FP aggregated in the cis leaflet such that lipids were depleted and replaced by peptides; even the formation of a peptide dimer is sufficient to exert this lipid-displacing effect (Fig. 5). The FP/lipid domain had increased water concentration in the bilayer midplane; such a thinner and more hydrated membrane domain is hypothesized to porate more easily than a peptide-free bilayer. Membrane elastic theory was extended to accommodate a new intermediate to water pore formation where a funnel-like defect, comprised of tilting FP, thins lipid further by a) moving the hydrophilic edge of the FP towards the membrane midpoint and b) minimizing the length of the hydrophobic defect that acts as the barrier to poration. Increasing FP density increased hydration in the MD-simulated lipid tail region of the trans leaflet, in agreement with theory predictions for greater water entry under a FP aggregate due to a shorter hydrophobic defect. Thus, the synergy of these two effects on the membrane hydrophobic region under a FP aggregate (FP tilting to thin and increased lipid hydration) can explain how FP facilitate water pores in bilayers at the critical time when fluorescence evidence for FP aggregation is detected.

The simulations presented here were not of sufficient length to observe pore formation and thereby directly test the analytical model presented. However, subsequent simulations of the 10 FP system with added lysolipids to destabilize the bilayer led to pore formation (manuscript in preparation); in these systems, initiation of pores involves a peptide tilt like that depicted in Fig. 8C (see Fig. S8 for definitions).

In the presence of FP, spectroscopic measurements demonstrated release of contents from large unilamellar vesicles (LUVs) composed of PC and PS/PC (1:1) ^27,45,46^, consistent with subsequent electrophysiological recordings ^23^, and electron and light microscopy ^22,25^. While these findings indicated that the influenza FP and its analogs destabilize the membrane, they did not address whether these permeability changes are associated with stable pores. Here, in single FP-GUV experiments, fluorescent dye influx continued over minutes, suggesting that the FP-induced influx does not proceed via a short lived transient flickering pore but rather via either a stable or constantly flickering pore ^33^. The lipid dependence of FP induced stable pore formation at neutral pH was the same as determined in the previous studies on intact virion and VLP ^25^.

Molecular dynamics simulations have demonstrated the ability of isolated FPs to distort the bilayer ^47^. These simulation results are consistent with several studies describing the interaction between the FPs and model membranes. For example, x-ray diffraction and differential scanning calorimetry indicate that the FPs destabilize DOPE membranes ^28,48^. However, the effect of cholesterol on the estimated depth of penetration of FP, based on trp fluorescence in vesicles containing singly brominated lipids is less than reported here ^49^.

MD simulations at higher density (6 and 10 FP) were motivated by the observation that hemagglutinin is densely packed on the virus surface. Specifically, the approximately 9.5 nm spacing ^22,50^ implies that a centered hexagon of 7 HA can be contained in a square patch of 15.3 nm/side. Since there are 3 FP per HA, in principle, 21 FP could maximally fit in such a patch; our 10 FP systems, while dense, are well below this upper limit. MD simulations of spontaneously emerging aggregates of FP on a single leaflet yielded qualitatively different results from those obtained at infinite dilution. Now, the FP aggregated and displaced many lipids in the cis (top) leaflet, thereby thinning that leaflet. Whereas a *single* FP coexists with a large ensemble of lipid configurations featuring tails that can spread under the FP, lipid tails adjacent to the FP *aggregate* were only partially able to shield the hydrophobic underside of that aggregate. Furthermore, at the highest FP densities (10 FP) the *trans* leaflet thins and becomes negatively curved (concave) as the terminal methyl groups interact with the remaining exposed regions of the aggregate’s hydrophobic underside (Figs. 5, S3, and S5). This curved region of the membrane is more permeable to water, as indicated by the substantial changes to the water PMF in the region of the FP aggregate and the increase in the water permeability (Fig 5). These structural changes to the bilayer are expected to modulate the spontaneous curvatures of the individual leaflets in ways not possible for non-interacting FPs.

Pores were not formed in any of the simulations, most likely due to an energy barrier incompatible with the microsecond atomistic simulation durations currently practical. Consequently, the analytic theory was developed to bridge the gap between simulation and experiments. Since the underlying ultrastructure of a FP membrane microdomain is radically different from that of a fluid mosaic FP-embedded bilayer, purely lipid leaflet elastic theory was modified to consider the work of bending the membrane into three possible intermediates of membrane poration. The tilting of FPs towards each other amid an aggregate, bringing their hydrophilic faces closer to the bilayer normal, was the most energetically favorable, with their hydrophobic faces shielding the largest fraction of the lipid tail from the full hydration of a water pore (see Fig. 8c). The use of membrane physical parameters of cholesterol or DOG-containing membranes in the calculations made pore formation less energetically favorable due to their significant negative MSC. Analytical calculations supported the experimental observation that the DOG-containing vesicles with the same negative MSC as cholesterol-containing ones exhibited a similar inhibition of poration (Fig. 1c), suggesting that it is not the chemical structure of cholesterol *per se* that is responsible for cholesterol’s pore neutralization activity. Compared to previous estimates for the lowest energy pathway in lipid bilayers composed of longer chain lipids, the FP/lipid hybrid bilayer has an even lower energy pathway to membrane poration^39,51-54^. Illya and Deserno observed qualitatively similar aggregation in their simulations of highly coarse grained peptide/bilayer systems ^55^; coarse grained methods should be useful for further studying aggregation of these FPs.

If an underlying evolved activity of the highly conserved FP is to aggregate, as demonstrated in this study and by the rosette formation described in the introduction, then aggregation may be due both to the hydrogen bond interactions of FP amino acid R groups seen in the MD simulations and membrane-mediated attraction shown in this work. Simulations of other amphipathic alpha helices^37^ do not show the aggregation described in this manuscript, potentially due to differential embedding of the two component alpha helices unique to FP. The deep embedding of the N-terminal helix, consistent with previous measurements by the Lentz lab, is striking ^49^. Is this aggregation of FP relevant to the membrane fusion of infection? A pore in a single bilayer (a target cell membrane) is not a fusion pore that links two bilayers (viral envelope and target cell membrane). Perhaps the work done by the FP microdomain comes later in the fusion process, after hemifusion diaphragm (HD) formation, when it would coat the merged leaflets trans to the HD. There it may induce a *bone-fide* fusion pore at the edge of the HD. In support of this idea, protein domain insertion at a distance (into a leaflet trans to the remodeling leaflet) lowers the activation barrier for fusion in two other membrane remodeling systems: PH domain insertion in dynamin-mediated membrane fission ^56,57^ and cell-cell fusion where myomaker acts after myomerger to form the fusion pore ^58,59^. In both systems, the protein domain action is thought to occur near or at a three-way lipidic junction ^56,58^. Alternatively, FP from intact enveloped virions might rupture endosomes and enter target cell cytoplasm for subsequent envelope disruption and intracellular trafficking of the viral genome to the nucleus. Regardless, the FP condensate revealed here, displacing lipids from one leaflet, is remarkably like the recent caveolin1 structure, also proposed to displace lipids from one leaflet to form a faceted surface of a caveolae^60^; here alpha-helices also stack and provide sufficient thickness to match the lipid leaflet thickness at the lipid/protein boundary.

## Materials and Methods

1-palmitoyl-2-oleoyl-*sn*-glycero-3-phosphocholine (POPC), cholesterol, and 1-2-dioleoyl-*sn*-glycerol (DOG) were from Avanti Polar Lipids, Inc. (Alabaster, AL) and other chemicals were from Sigma-Aldrich (St. Louis, MO), unless specified. Stock solutions of POPC (13.15 mM), DOG (3.22 mM), and cholesterol (20 mM) were made in CHCl_3_ (Burdick & Jackson, Mexico City, Mexico, high purity solvent) and stored in brown glass vials at –20 °C. Structures of FP analogs (synthetic amphipathic peptide with additional hydrophilic amino acids to their C-terminus) in the presence of phospholipid bilayer membranes or micelles are alpha-helical in their secondary structure with either a closed coiled-coil tertiary structure ^61^ or an open ‘boomerang’ shaped tertiary structure^62^. A 23 amino acid synthetic peptide corresponding to the X-31 FP domain^56^ was custom synthesized by GenScript, NJ labelled with tetra-methyl rhodamine (Sequence: GLFGAIAGFIENGWEGMIDGWYG [LYS(TRITC)]). Stock solutions of FP in DMSO were stored at – 20 °C. To compare the FP-induced membrane poration with intact X-31 influenza virus-induced poration ^10^, this X-31 FP sequence was used^56^. To compare poration results to MD simulations, the H-serotype FP sequence was used^55^ (GLFGAIAGFIEGGWTGMIDGWYG). Poration results were similar for both FP sequences, irrespective of dye labelling.

### Experimental Conditions

#### Preparation of Giant Unilamellar Vesicles

GUV were prepared by the gel swelling method as described previously ^63^. Briefly, to prepare GUVs of a specific lipid composition the required amount of lipid drawn from individual stock solutions was diluted into 200 µl of CHCl_3_ to a concentration of 3.94 mM (3.35 mg/ml). The mixture was vortexed with ∼10 µl of MeOH for ∼2 min to avoid incomplete mixing and obtain a clear solution in CHCl_3_. 10 µl of DiD (Invitrogen; prepared in DMSO as a 5 µM stock solution) was added to this lipid mixture and vortexed for ∼2 min. The lipid mixture (in CHCl_3_) was then deposited on a plasma cleaned (using a Harrick plasma cleaner, Ithaca, NY) microscope cover glass, coated with 5 % (w/w in ddH_2_O) polyvinyl alcohol (Merck Millipore). The organic solvent was evaporated by a gentle stream of nitrogen and then stored in high vacuum for 1 hour. The lipid film containing cover glass was then transferred to a 30-mm tissue culture dish. 500 µl of GUV formation buffer (1 mM EDTA, 1 mM HEDTA, 10 mM PIPES, 100 mM KCl, pH 7.4 with ∼200 mM sucrose) was added covering the entire surface of the cover glass and allowed to incubate for 30 min in the dark. After 30 min, the GUVs were harvested by gently tapping the sides of the dish, then gently removing the GUV suspension using a 1 mL pipette without touching the surface and transferring to a 1.5 ml micro-centrifuge tube. The GUV suspension was stored at 4 °C until further use. The total lipid concentration of the GUV suspension was 1.35 mg/ml (1.58 mM). Typically, GUVs were made the same day of the experiment. To vary the MSC systematically, POPC GUVs of varying concentrations of cholesterol, or DOG were prepared.

#### GUV poration assay

FP was added from the stock solution to a suspension of GUVs at a peptide to lipid ratio of 1:200 in a micro-centrifuge-tube. The total volume was diluted to 500 µl containing Alexa 488 or SRB. The mixture was then incubated at 37 °C for 15 min. While the FP-GUV mixture was incubating, 500 µl of a 5 mg/ml β-casein solution was added to a delta-TPG 0.17 mm dish (Bioptechs, Butler, PA) allowed to sit for 10 min to passivate the glass surface, and then washed 10 times with ddH_2_O. The FP- GUV suspension was then vortexed for 30 s and transferred to this imaging dish. For a given lipid composition, ∼50 - 100 GUV were observed and scored based on whether they have undergone influx of Alexa 488. Leakage was determined ∼15-18 min after FP addition to ensure equilibrium. The fluorescence intensity within GUV was monitored for an additional 30 min; no changes in the mean fluorescence intensity were detected after the additional waiting period. For kinetics measurements, the FP was directly added to the chamber contacting GUVs while mounted on the microscope objective.

#### Confocal microscopy

GUVs were imaged on a Zeiss LSM 880 microscope using a 63× oil 1.4 NA Plan-Apochromat objective. The objective was heated (Bioptechs, Butler, PA) to maintain 37 °C inside the chamber (verified with a microprobe thermistor). Alexa 488, SRB and DiD were excited with 488, 543 and 633 nm lasers respectively and detected using PMT detectors on separate imaging tracks.

#### Image, data, statistical and aggregation analyses

Microscopy images were analyzed using ImageJ (NIH). Data were analyzed using SigmaPlot (Systat Software, Inc., Chicago, IL), Excel (Microsoft, Inc., Redmond, WA), and MATLAB (The MathWorks, Inc., Natick, MA). The aggregation hypothesis predicts higher than expected concentrations of bound FP during poration. The polarization properties of a high NA objective produce a cosine-squared dependent modulation of an otherwise constant intensity ^64-66^. To correct the intensity profiles recorded on GUVs, template matching alignment was employed (to minimize position jitter noise), using the ImageJ plugins (Template Matching and Oval Profile Plot). However, neither the observed profiles at a single time point, nor averaged over a subset of time points (5, for example) nor the entire time series could be described using either a single uniform orientation or two component mixtures of orientations, unlike the profiles predicted for a chromophore with identical orientations uniformly bound to the surface. Statistical evidence of FP aggregation on GUV includes pixels with larger than expected intensity, requiring consideration of both the slowly varying modulation around the circumference and the magnitude of the intensity, since variance increases with intensity. The aggregation analysis first calculated the intensity surface at the time of poration followed by an analysis of the distributional properties of the residuals. Direct calibration issues were avoided by normalizing the GUV pixel intensities into units of solution intensity calculated from the distribution of intensities along a 1-pixel wide ring matching the size of a GUV in cross-section (Fig. 9, inset). Since the distributions of FP intensity in solution, over time, often captured transient, FP-bound membrane structures, the distributions were modeled as a two component Gaussian mixture, with the lower intensity mean taken as the FP intensity in solution to be used in the normalization of membrane-bound FP. Possible changes to the chromophore properties following FP binding to the membrane were not observed because the initial normalized GUV intensity is approximately 1 (Figure 8; solid horizontal line at ∼ 1 for normalized GUV intensity values less than t_*crit*_ = 147 sec). Consequently, the intensity on the GUV surface is expressed in units of solution intensity that is directly related to the FP concentrations and designated as the normalized FP density. Note, the solution values are invariant with time, whereas the intensity of the FP bound to the GUV begins to diverge and increase with time. The extent of Alexa 488 influx (*R* = *F*_*in*_/*F*_*out*_; green trace) is also plotted and the time of poration indicated by the red arrow.

**Fig. 9.**
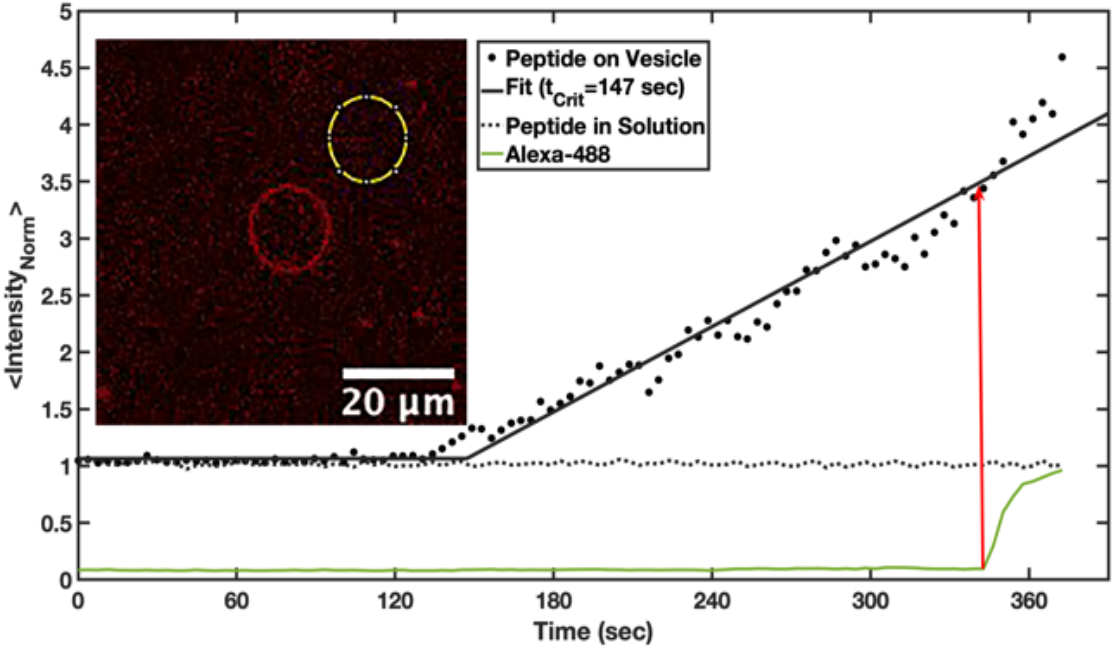
Kinetics of poration and FP accumulation on a POPC GUV. FP fluorescence signals were normalized by the FP mean intensity in solution. The time FP begins to accumulate on the vesicle, t_*Crit*_, was determined using piecewise linear fitting. The time, t_*Pore*_, poration begins (red arrow) was determined using piecewise linear and exponential fitting. In this example, the characteristic time between FP accumulation and poration is 197 sec with a mean FP intensity at poration 3.5 times the solution intensity. Inset) Confocal image of an FP labeled vesicle (red) and control ring (yellow), positioned in the solution, used to evaluate circumferential intensity of both the vesicle and solution as a function of time.

A window +/-2 timepoints around the estimated poration time, t_Pore_, was analyzed for the presence of pixels having FP intensity larger and smaller than expected. The data surface, defined by 3 parameters (radial angle, time, pixel intensity) were fit using locally weighted, smoothing quadratic regression (Lowess quadratic) with a data span of 20% to capture the complexity of the time, angle, and intensity of the peptide on the GUV. The same surface fits in solution were approximately planar. The spline-smoothed circumferential mean (over the five time points) removed any large scale (quadrant angle level) polarization-dependent intensity modulation of vesicle-bound FP. The resulting intensity deviations (residuals) were compared to the distributional behavior of FP in solution. The residuals represent both the positive and negative deviations around the circumferential mean; larger (smaller) than expected intensity deviations on the vesicle, when compared to the solution distribution are evidence for pixels with greater (lower) FP density. The spatial distribution of the FP intensities in solution represents the expected variation for orientationally random and spatially uniformly distributed data since the surface was a) approximated by a plane at a constant value of 1 (the normalized intensity) and b) the residuals were symmetric around 0.

### Molecular Dynamics Simulations

#### System preparation

Two membrane compositions were compared: pure POPC and 1:1 chol:POPC. Large systems were constructed with 10, 6, or 1 copies of the 23-residue NMR structure of the FP (sequence: GLFGAIAGFIEGGWTGMIDGWYG) determined by Lorieau et al.^61^ placed in the cis (top) leaflet of ∼150×150 Å membranes. Smaller symmetric systems were also constructed with a single copy of the FP in each leaflet of ∼75×75 Å bilayers for the estimates of spontaneous curvature listed in Table S1. All systems were constructed using the CHARMM-GUI *Membrane Builder* ^67-70^ with a water thickness of 17.5 Å and 150 mM potassium chloride in the solvent. Simulations utilized the CHARMM36 force field ^71,72^ and TIP3P water model ^73,74^, and were all performed in the isothermal-isobaric (NPT) ensemble with a 2.0 fs time step.

#### Long time scale simulations

All large simulations were simulated first in OpenMM version 7.4.1 ^75^ using the Rickflow package version 0.7.0. OpenMM equilibration times were 1000 ns for 10 FP systems and 100 ns each for 6 and 1 FP systems. Following this, because of the large fraction of FPs in the cis leaflet, the 10 FP systems were simulated for 30 ns in CHARMM ^76^ using P2_1_ boundary conditions ^77^ to allow lipids to equilibrate between leaflets. The number of lipids per leaflet was averaged over the last 7.5 ns of these P2_1_ simulations; representative frames with lipid distributions matching this average were selected as starting coordinates for simulation on Anton 2 ^78^. All large systems were then simulated on Anton 2 for production runs. The Anton 2 simulations times were 20 μs for 10 FP systems, and 2 μs each for 6 and 1 FP systems. The symmetric single FP systems were simulated for 500 ns in OpenMM. As controls, both system sizes, FP-free systems of comparable size (150×150 Å and 75×75 Å) were simulated for 500 ns in OpenMM

All OpenMM simulations were performed at 310 K with the Nose-Hoover chain ^79-81^ velocity Verlet integrator implemented in Openmmtools ^82^ and the Monte-Carlo membrane barostat ^83^. Bonds with hydrogen were constrained using the SETTLE and CCMA algorithm ^84,85^. A 12.0 Å cutoff was used, with a force-switching function from 8−12 Å. Long-range electrostatics were treated using the particle mesh Ewald method ^86^. Coordinates and velocities were saved every 50 ps. Simulations on Anton 2 utilized the Multigrator framework ^87^ with the Nose-Hoover thermostat (310 K) and semi-isotropic MTK barostat. An 8 Å real space cutoff distance was used, and long-range electrostatics were evaluated with the u-series method ^78^. Coordinates were saved every 200 ps.

#### Simulation analysis

Thickness maps and leaflet position maps were calculated using the MEMBPLUGIN ^88^ extension for VMD, and all system snapshots were rendering using VMD ^89^. Lipid density plots were calculated using LOOS ^90,91^. FP clustering was assessed using the cluster module of the *freud* Python library^92^. All FP heavy atoms were considered in the analysis, and a cutoff of 3.5 Å, consistent with a typical hydrogen bond donor-acceptor cutoff distance, was used to define points as belonging to the same cluster. Peptide tilt angles were calculated by defining a tilt vector between the alpha carbon of residue 13 (midpoint of the bend) and the geometric center of the alpha carbons of residues 3 and 3 (the helical residues furthest from the bend) and calculating the angle of this vector with respect to the xy-plane. Roll angles were calculated in a similar fashion, with the roll vector defined between the alpha carbons of residues 3 and 22 (see sketch in Fig S8). Water, lipid, and peptide density distribution profiles were calculated using the density function in *cpptraj* ^93^ and a 0.5 Å bin width; only atoms within a cylinder of 30 Å centered about the origin (Fig. S7) in the xy-plane were considered for this calculation (FP-containing systems were first imaged such that the FP aggregate was centered at the origin, in the plane of the membrane.) The PMF, *F*(z), was calculated from the normalized water density distribution, *p*(z):

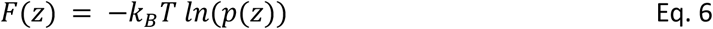

#### Permeability and partition coefficients

The inhomogeneous solubility diffusion (ISD) model ^35^ provides a simple relation between the free energy profile and the resistance to permeability, *R*. The permeability *P* in the ISD model is written:

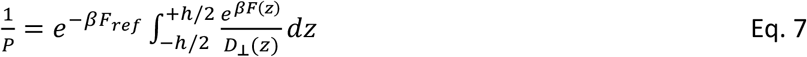

where *D*_⊥_(*z)* is the position dependent diffusion coefficient, *β* = 1/*k*_*B*_*T, h* is the thickness of the bilayer, and *F*_*ref*_ is the reference free energy in the water phase that is set to zero. The diffusion constant is not calculated here and will be assumed to be a constant ⟨*D*⟩. The resistance is then defined from Eq. 4 as:

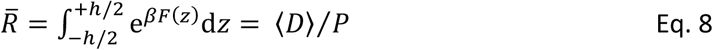

Note that the preceding definition of 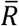 differs slightly from the usual definition of *R*, where the diffusion constant remains in the integral.

### Calculations of the energy of elastic deformations of lipid membranes

To calculate the energy of membrane deformations, we utilize the Hamm and Kozlov model ^38^ according to which the lipid monolayer is considered to be an elastic continuous 2D media. The state of the lipid leaflet in this model is defined by the field of unit vectors **n** corresponding to the average direction of the lipid molecules, and the shape of the monolayer-water interface. We assumed that the system has an axial symmetry with respect to the axis passing through the center of the FP cluster or single FP. The energy functional comprises four contributions: bending (that is defined by the mean curvature *J*), tilt (**t**), lateral compression/stretching (*α*) and Gaussian curvature (*k*_*g*_) energy terms:

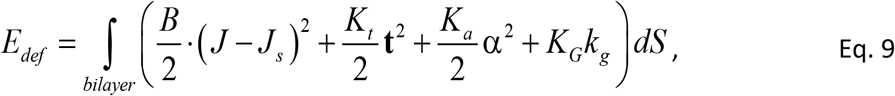

where *B, K*_*t*_, *K*_*a*_, *K*_*G*_, are bending, tilt, compression/stretching and Gaussian elastic moduli; *J*_*s*_ is the monolayer spontaneous curvature. The integration is performed over both monolayer surfaces. The leaflet’s effective curvature is determined by the divergence of director field: *J* = −div(**n**). The tilt vector is defined as a deviation of the lipid director from the normal **N** to the leaflet’s surface: **t** = **n** – **N**. The Gaussian curvature in the axially symmetric system is defined using the following product: 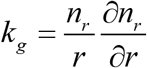, where *n*_*r*_ is the projection of the director on the O*r* axis. The following parameters were used to calculate the energy of membrane deformations: Hydrophobic thickness and lipid MSC (Table S1) and were obtained from MD simulations; *J*_*s,POPC*_ = –0.043 nm^−1^, *J*_*s,mix*_ = –0.173 nm^−1^ for pure POPC and mixed chol:POPC membrane leaflets; bending rigidity for pure POPC membrane *B*_POPC_ = 11.2 *k*_*B*_*T* ^94^ and for mixed chol:POPC *B*_*mix*_ = 13.7 *k*_*B*_*T* ^95^. The lateral area of FP was taken to be 4 nm^2^. We used the following values for the remaining elastic constants: *K*_*t*_ = 10 *k*_*B*_*T*/nm^2^, *K*_*a*_ = 30 *k*_*B*_*T*/nm^2^, *K*_*G*_ = –0.3·*B k*_*B*_*T* equal for both pure POPC and mixed membranes.

To calculate the elastic energy due to insertion of a single FP, *E*_*FP*_, fusion peptides and mini-cluster were considered as rigid objects that impose boundary conditions on the directors ^43,44^ *n*_*r*_ (*r* = *r*_0_, φ), *n*φ (*r* = *r*_0_, φ), where *r*_0_ is the radius of single FP or their cluster and *n*φ is the projection of the director on the *Oφ* axis; parameters were as above. Directors and leaflet surface were constrained to be continuous everywhere except in the regions occupied by the FPs. The membrane was required to be unperturbed far from the cluster boundary. The elastic energy was minimized with respect to the membrane deformations and FP’s insertion depth. The detailed description of the calculation procedure can be found in Kondrashov, et al. 2018 ^44^.

## Author Contributions

SH, EW, PSB and JZ designed and carried out the microscopy experiments. AR and RWP designed and carried out the MD simulations. SAA and TRG developed the analytical theory. All authors analyzed the results and wrote the paper.

## Acknowledgments

This work was supported by the Intramural Programs of the *Eunice Kennedy Shriver* National Institute of Child Health and Human Development and the National Heart Lung and Blood Institute of the NIH, using NIH high performance computational resources (NHLBI LoBoS and HPC Biowulf cluster). Anton 2 computer time was provided by the Pittsburgh Supercomputing Center (PSC) through Grant *R01GM116961* from the NIH; the Anton 2 machine at PSC was generously made available by D.E. Shaw Research. TRG is grateful to the Russian Science Foundation (Project No. 19-74-00152) for support. We thank Dr. M. Garten for helpful discussion regarding GUV preparation and Dr. A. Bax for a kind gift of an FP analog for a control experiment.

## Data availability statement

All data generated and analyzed in this article and its supplementary information files are available from the corresponding author on reasonable request.

