## Supplemental Information for "Planar aggregation of the influenza viral fusion peptide alters membrane structure and hydration, promoting poration"

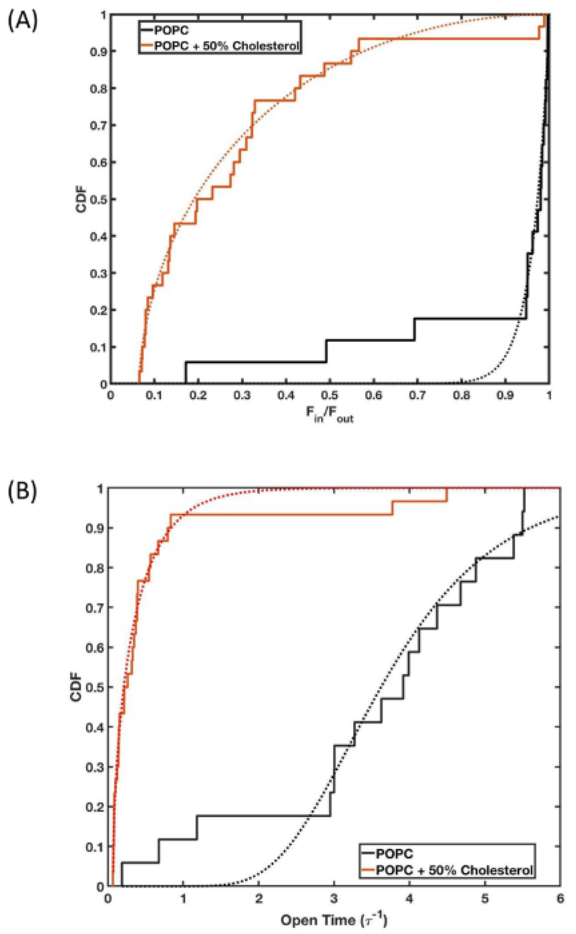


**Fig. S1.** **Effect of cholesterol on fusion peptide induced pores.** (A) Cumulative distribution functions (CDF) of the influx ratio ($R = F_{in}/F_{out}$) for POPC and POPC + 50% cholesterol GUV are described by generalized Kumaraswamy distributions[1]: (CDF = $1 - (1 - z^{\alpha})^{\beta}$; $z = (R - R_{\min})/(1 - R_{\min})$), with distribution shape parameters $\alpha$ and $\beta$, and $R_{\min}$ = 0 for pure POPC and $R_{\min}$ a fitting parameter for POPC and 50% cholesterol [1,2]. For POPC GUV, $\alpha$ = 24.0 (15.1, 32.8) and $\beta$ = 0.94 (0.64, 1.24) while in the presence of 50% cholesterol $\alpha$ = 0.59 (0.52, 0.66) and $\beta$ = 1.91 (1.61, 2.21) with $R_{\min}$ = 0.065 (0.063, 0.067) (fit values (95% lower, upper confidences)); these parameter sets correspond to fractional filling distributional means of 0.96 and 0.26 or median quantiles of 0.97 and 0.13 for POPC in the absence and presence of 50% cholesterol, respectively. (B) Cumulative distribution functions for the influx time for POPC in the presence and absence of cholesterol following remapping of R onto the general kinetic relationship $R = (1 - e^{-t/\tau})$ where $\tau$ is a characteristic time constant for the influx process. The cumulative distribution functions for the open time for POPC, as a function of time in units of $\tau$, in the presence and absence of cholesterol are described by generalized exponential distributions (CDF = $(1 - e^{(-\lambda(t/\tau - (t/\tau)_{\min}))})^{\alpha}$) with location ($(t/\tau)_{\min}$), shape ($\alpha$), and scale ($1/\lambda$) parameters.[3]

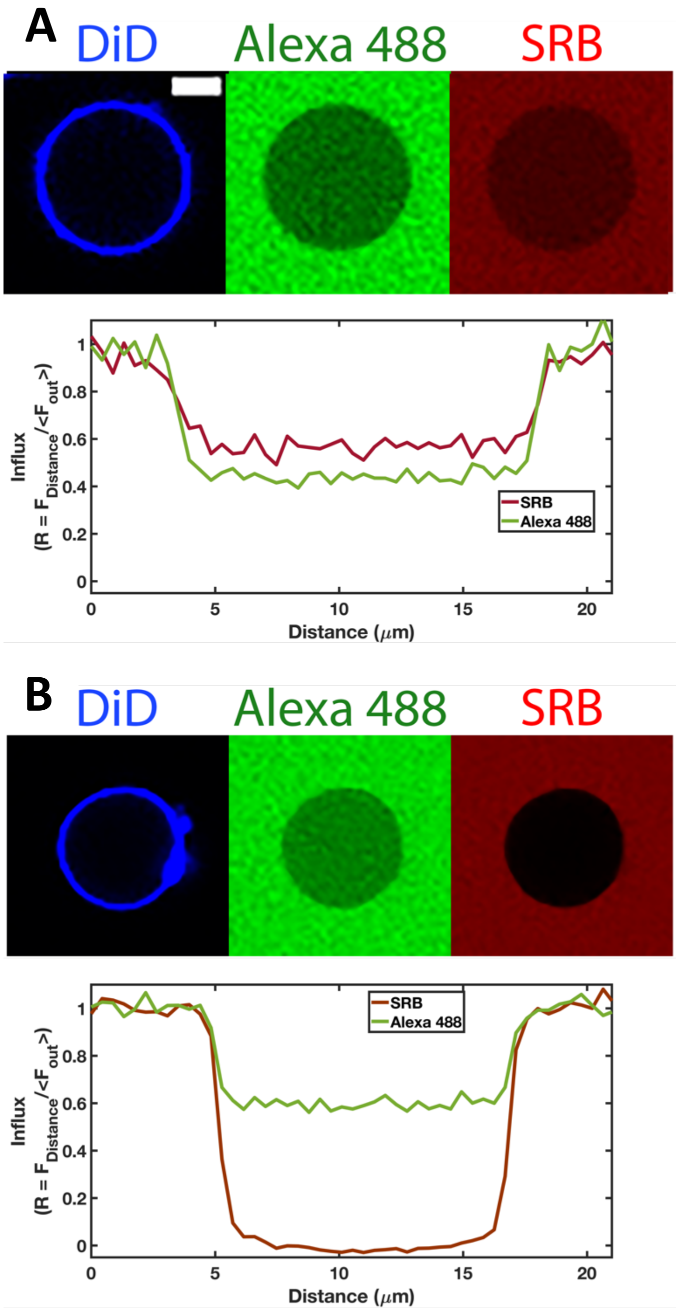


**Fig. S2. Closure of fusion peptide-induced pores in the presence of cholesterol**. (A) Similar degrees of influx were observed when Alexa 488 (green) and SRB (red) were introduced simultaneously. Equatorial line profiles of Alexa 488 (green) and SRB (red) quantitatively shows influx. GUVs containing 50 mol % cholesterol were incubated with Alexa 488, and SRB and FP were added to the system. (B) Restricted dye entry and exclusion of second dye from vesicles. FP was introduced to a suspension of GUVs in the presence of Alexa 488. After 25 min SRB was added to the same suspension of GUVs. Note the differential extent of the influx. Intensity of SRB inside the GUV is negligible compared to Alexa 488. Scale bar is 10 μm.

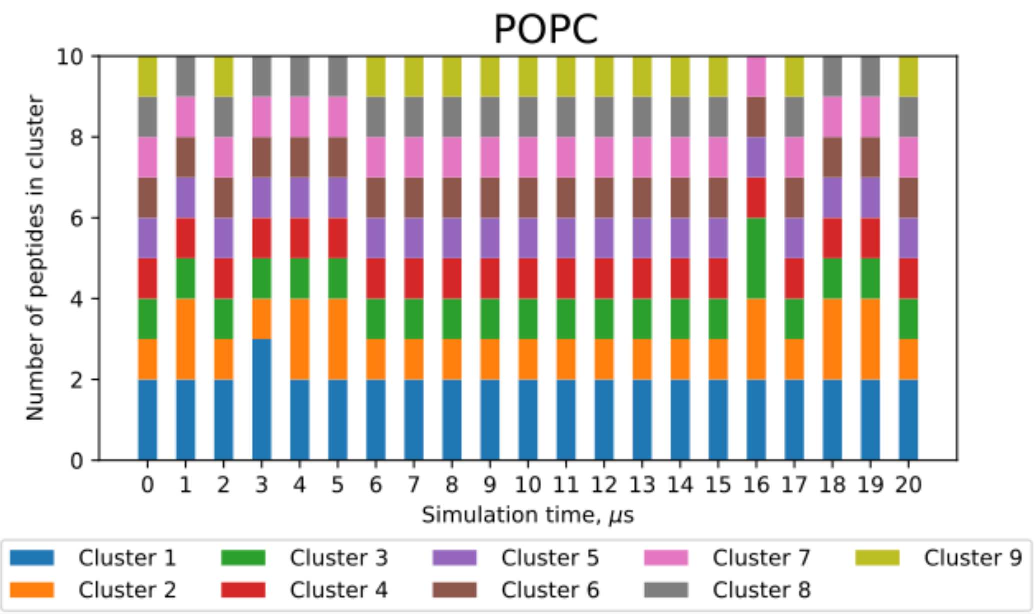


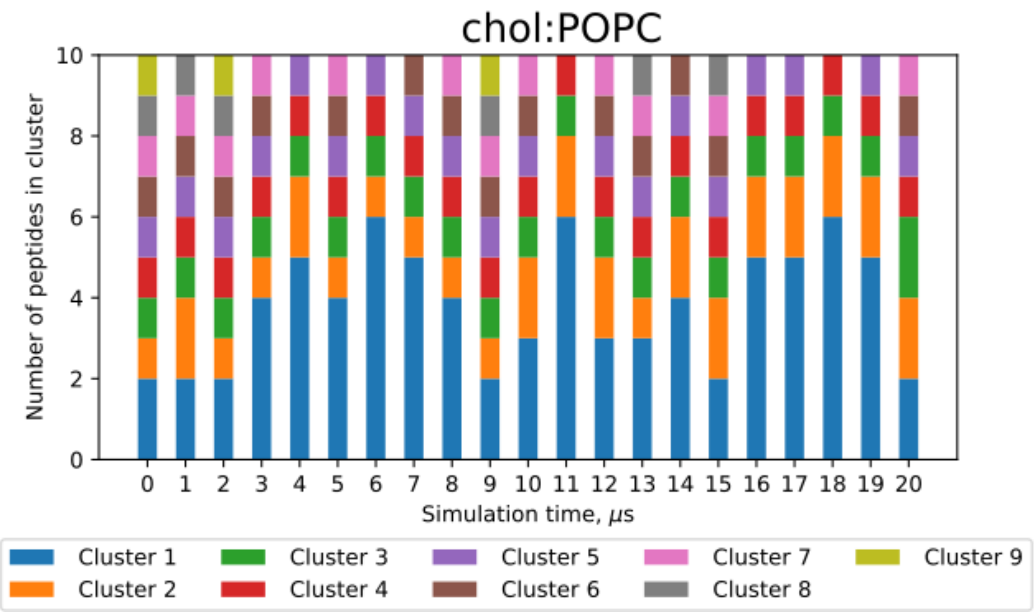


**Fig. S3. Clustering in 10 FP systems**. Clustering shown every microsecond of the 20 μs production simulation. The number of peptides in each cluster is denoted by the different colors in the bar graph.

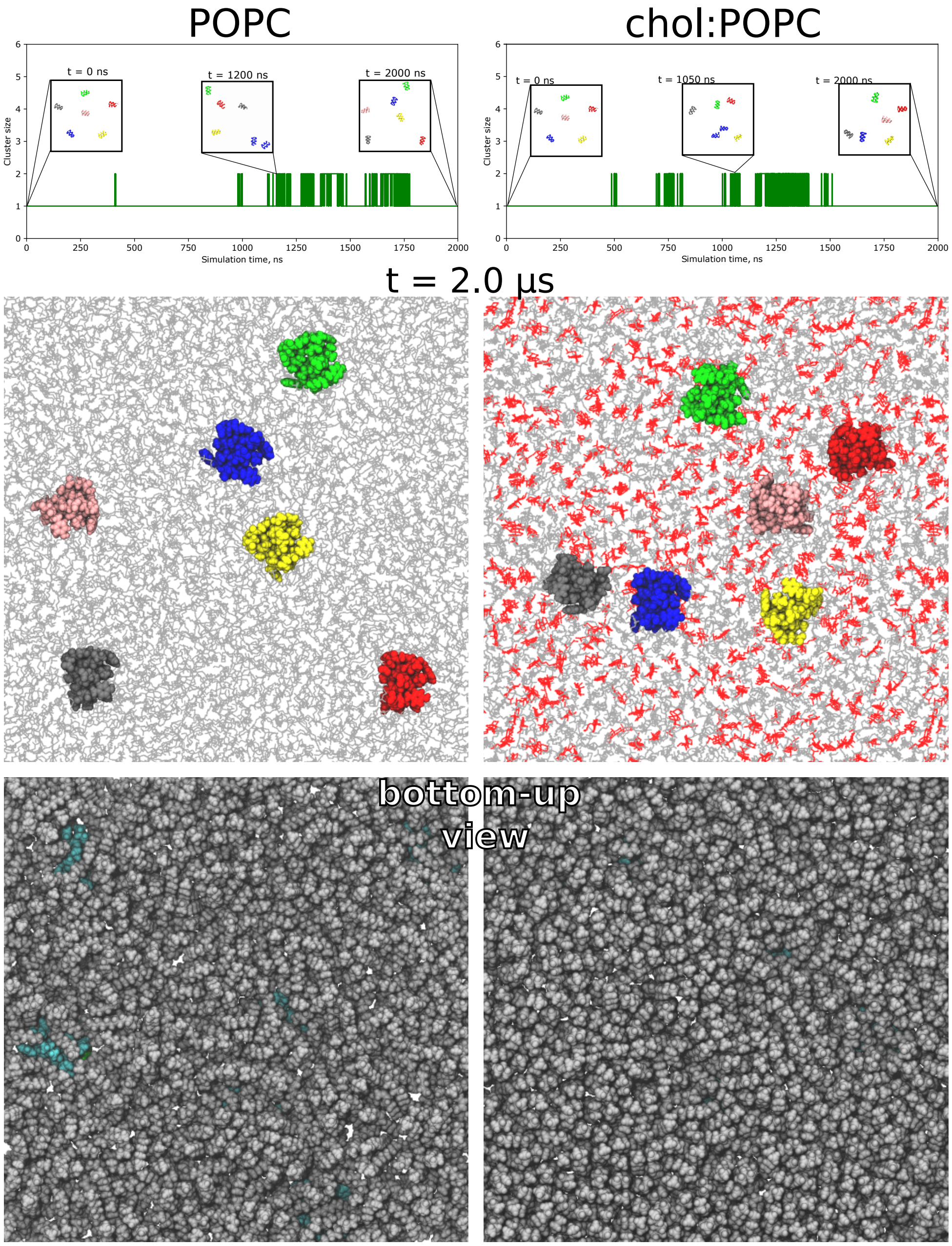


**Fig. S4. Peptide clustering is limited in 6 FP simulations**. Top) Largest cluster size as a function of simulation time for 6FP:POPC (left) and 6FP:chol:POPC (right). Insets show example snapshots. Middle) Final peptide locations after 2000 ns in POPC (left) and chol:POPC (right). Peptides are shown in a VdW sphere representation which each peptide in a different color. POPC and cholesterol are shown in grey and red line representations, respectively. Bottom) Bottom-up view of the top leaflet from the snapshots shown at center. The hydrophobic undersides of the peptides (cyan VdW sphere representations) are mostly covered by the hydrophobic lipid tails (grey).

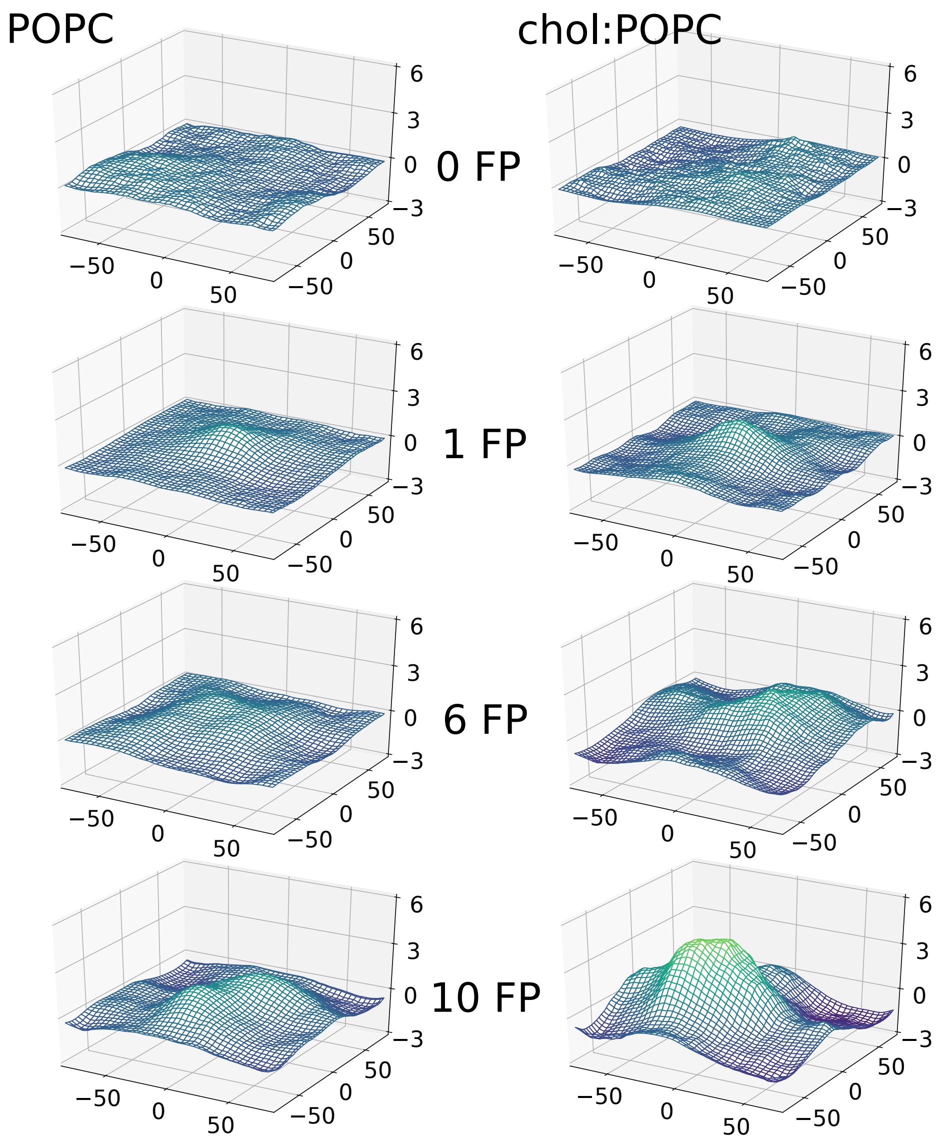


**Fig. S5. Trans leaflet curvature maps for all simulated systems.** Trans leaflet surface plots for the POPC (left) and chol:POPC (right) systems, peptide-free or in the presence of 1, 6, or 10 FP in the cis leaflet. All surface plots are calculated over the final 800 ns of each trajectory, except for the peptide-free systems which utilized the full 500 ns trajectory. Z = 0 has been set to the average position of all POPC C2 atoms in the leaflet. Note that the scale in z differs from that of the xy-plane for greater clarity. All axis labels are in Å.

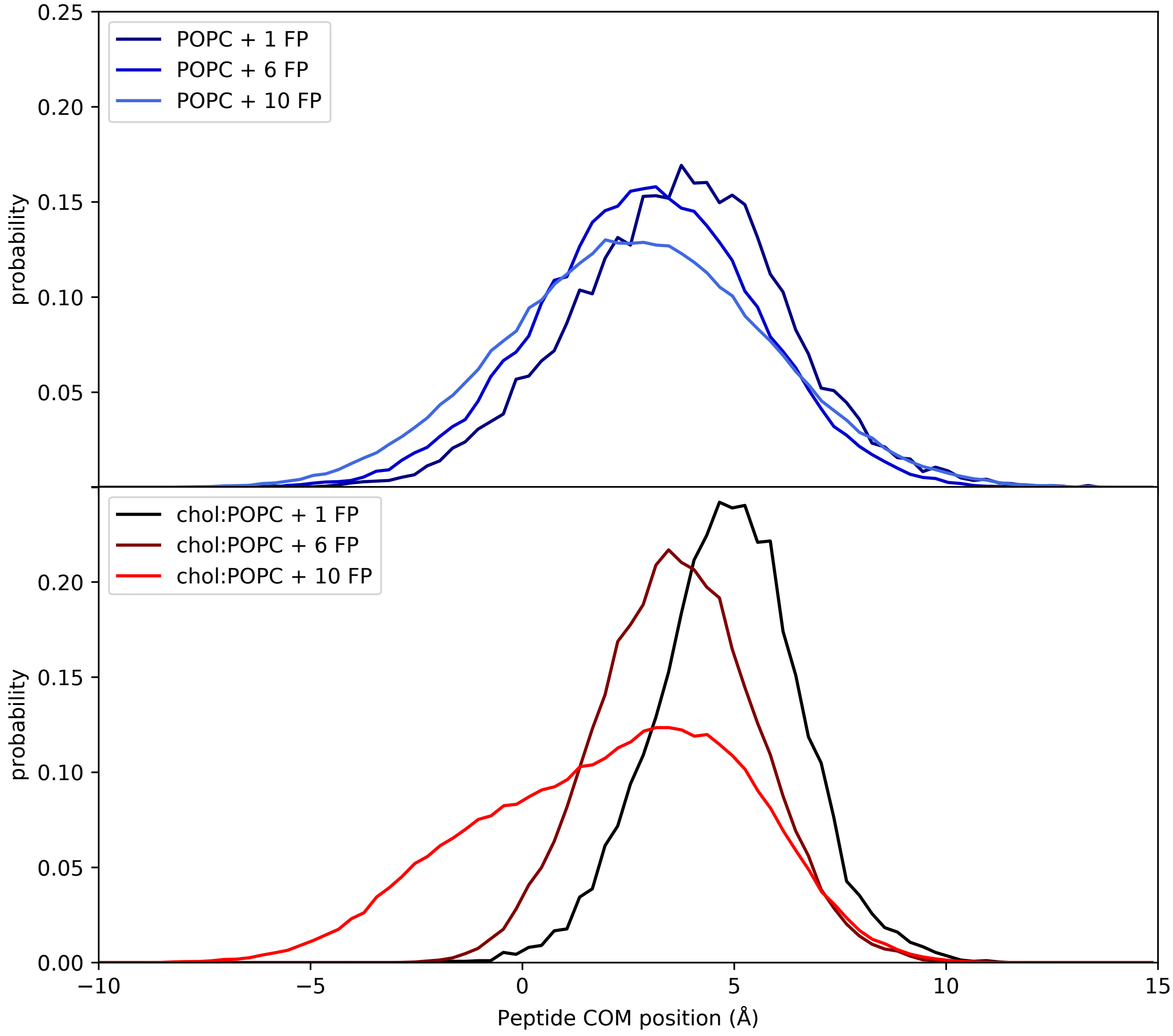


**Fig. S6. Peptide center of mass location**. Probability distributions of the peptide center of mass location with respect to the POPC C2 plane for the pure POPC (top panel) and chol:POPC (bottom panel) systems. The peptide aggregates insert more deeply than the isolated peptide, regardless of the bilayer composition.

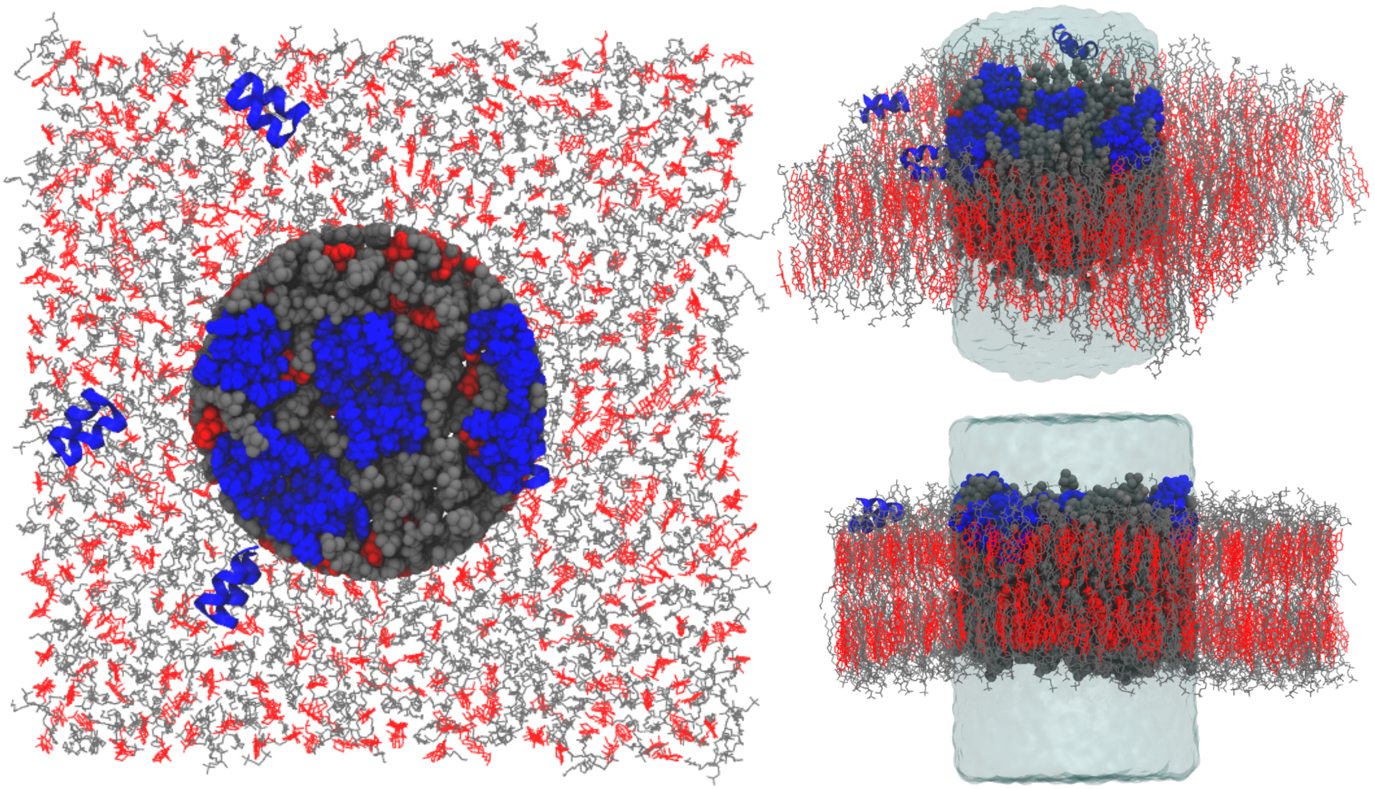

**Fig. S7. Visualization of the cylindrical region used for density profile calculations**. Molecules within the cylinder, with radius 30 Å centered at the origin, are depicted in a space-filling representation, while the water within the cylinder is depicted as a transparent bubble. POPC and cholesterol are shown in grey and red, respectively; peptides are in blue; water outside the cylinder is not shown.

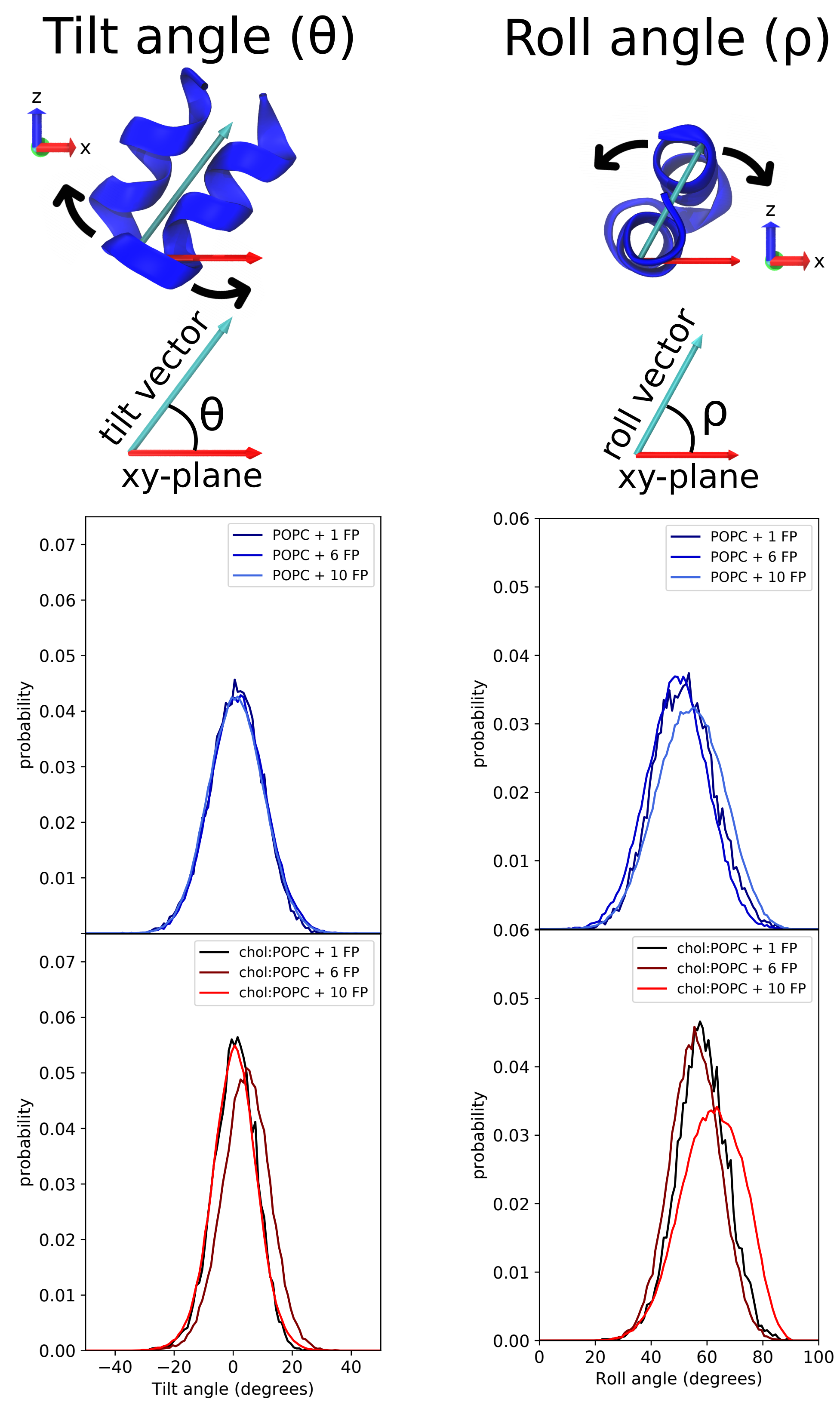


**Fig. S8. Tilt and roll angles for all eight systems simulated.** Peptide tilt and roll are defined in the top panel. While there is no difference in the tilt angle between peptides in POPC compared to 1:1 chol:POPC, the average roll angle is larger by 4.5 degrees in the 1:1 chol:POPC membrane.

| System | $R_0$ (Å) | $C_0$ (Å$^{-1}$) | F'(0) | $K_{c,m}$ (kcal/mol) |
|---|---|---|---|---|
| POPC | -235 | -0.0043±0.0008 | 0.0407±0.0072 | 9.55[a] |
| POPC:2FP | -297 | -0.0034±0.0013 | 0.0322±0.0122 | 9.55[a] |
| chol:POPC | -58 | -0.0173±0.0046 | 0.2018±0.0542 | 11.67[b] |
| chol:POPC:2FP | -129 | -0.0078±0.0026 | 0.0907±0.0299 | 11.67[b] |

**Table S1. Spontaneous curvature generation by a single FP per leaflet.** The radius of curvature, $R_0$, is calculated as $1/c_0$. The spontaneous curvature is calculated from *F'(0)*, the derivative of the leaflet's free energy with respect to curvature, which is computed from the first moment of the lateral pressure profile. [a]Caclulated from bilayer bending constant reported by Venable et al.[4]; [b] Estimated using POPC $K_{c,m}$ and data from Chen and Rand[5].

**Videos S1-S6. Movies of final frames of all 6 MD simulations with FP depicting representative FP orientations.** Peptide backbone residues are shown in ribbon diagrams, with each FP colored a different color. POPC phosphorus atoms are depicted as gold spheres to orient the viewer. All other atoms, including lipids, water, and protein side chains, are omitted for clarity. The simulation is oriented with the membrane normal "up", and in the video the view rotates twice (720°) about the membrane normal.

### Appendix: Cluster Mechanism

We consider the system of $N$ FPs occupying area fraction $x$ of one leaflet of a lipid bilayer membrane. The energy of the $N$ isolated FPs equals $N \cdot E_{FP}$, where $E_{FP}$ is the energy due to insertion of a single FP (see Material and Methods). Since FPs replace lipids from the cluster, the energy of the cluster of radius $R$ is stored in its boundary. Neglecting direct FP-FP interactions, $E_{cluster} = 2\pi R\gamma(R)$, where γ is the cluster line tension. The dependence of the cluster line tension on its radius is weak (Fig. S9).

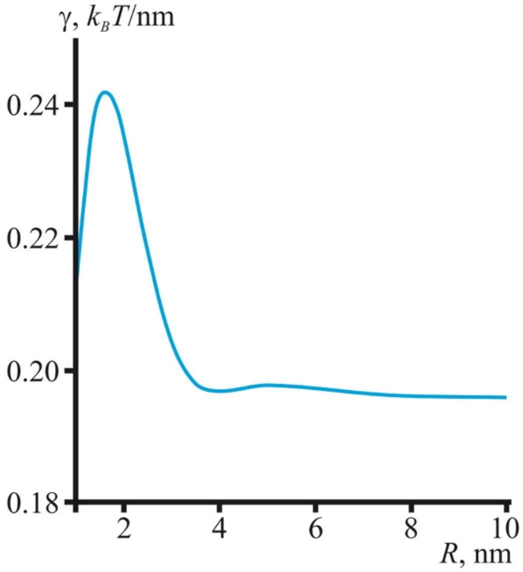


Fig. S9. Dependence of the cluster line tension ($k_BT$/nm) on its radius. FP monolayer thickness is taken as $h_{FP}$ = 1.1 nm, whereas the surrounding membrane thickness $h$ = 1.5 nm.

The radius of the cluster is derived from the number of FPs $n_{clust}$ in it:

$$R = \sqrt{\frac{n_{clust} a_0}{\pi}}, \qquad \text{Eq. S1}$$

where $a_0$ is the area of a single FP. The entropy-related work of forming a cluster containing $n_{clust}$ FPs with mole fraction equal to 1 (no lipids inside the cluster) is

$$E_{ent} = k_B T\left(N\ln\left(\frac{1}{x}\right) - \frac{N}{n_{clust}}\log\left(\frac{n_{clust}}{x}\right)\right), \qquad \text{Eq. S2}$$

where $x$ is the area fraction of the FPs in the system. The total energy difference between the system of separated FPs and the ensemble of the clusters containing $n_{clust}$ FPs each is:

$$\frac{\Delta E}{N} = \frac{E_{cluster} - N \cdot E_{FP} + E_{ent}}{N} = 2\pi\sqrt{\frac{a_0}{\pi n_{clust}}}\gamma - E_{FP} + k_B T\left(\ln\left(\frac{1}{x}\right) - \frac{1}{n_{clust}}\log\left(\frac{n_{clust}}{x}\right)\right). \qquad \text{Eq. S3}$$

Clustering is favorable if the energy difference $\Delta E$ is negative, i.e. at concentrations $x$ above some critical value $x_c$, i.e. when

$$\Delta E(x = x_c) = 0. \qquad \text{Eq. S4}$$

For large clusters ($n_{clust}$ >> 1), we can neglect the first and the last terms in Eq. S3. This is equivalent to neglecting the cluster boundary energy and the entropy of the cluster ensemble, significantly simplifying Eq. S3 to

$$\frac{\Delta E}{N} \approx -E_{FP} + k_B T \ln\left(\frac{1}{x}\right) \qquad \text{Eq. S5}$$

This leads to the simple solution of Eq. S4 for the critical concentration of FP for cluster formation:

$$x > x_c^0 = e^{-\frac{E_{FP}}{k_B T}}. \qquad \text{Eq. S6}$$

The dependence of the energy difference $\Delta E$ on the FP concentration $x$ is shown in Fig. S10. The black point indicates the exact solution of Eq. S4, whereas the vertical red line indicates the solution for $x_c^0$ according to the approximate Eq. S6.

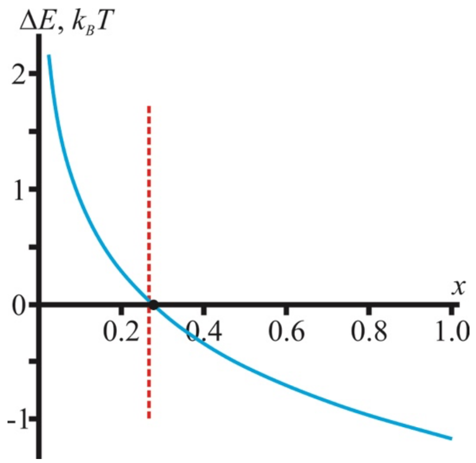


Fig. S10 Dependence of the energy difference $\Delta E$ on the FP concentration $x$. FP area $a_0$ = 3 nm$^2$, cluster size $n_{clust}$ = 10, isolated FP energy $E_{FP}$ = 1.3 $k_B T$. The vertical red line is the solution of Eq. S6 for $x_c^0$.

The dependence of the critical FP concentration $x$ on the cluster size appears to be very weak (see Fig. S11), and Eq. S6 gives a very good estimation for the critical concentration $x_c^0$.

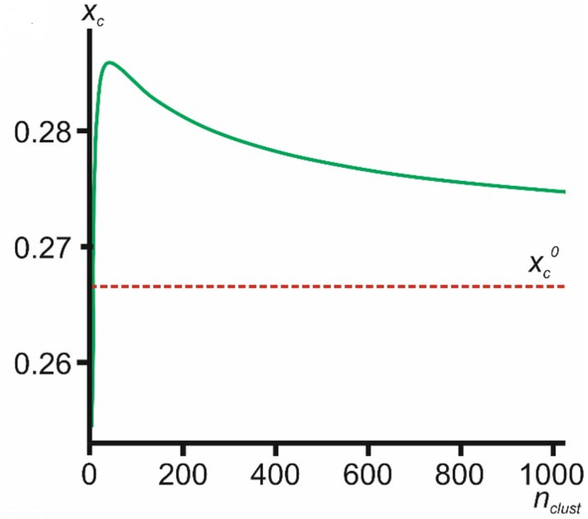


Fig. S11. The dependence of the critical FP concentration $x_c$ on the cluster size. The horizontal red line is the solution of Eq. S6 for $x_c^0$.
